# Targeting lysosomal damage is a new therapeutic perspective for Duchenne Muscular Dystrophy

**DOI:** 10.1101/2025.02.11.637610

**Authors:** Abbass Jaber, Laura Palmieri, Rania Bakour, Elise Lachiver, Ai Vu Hong, Nathalie Bourg, Jérôme Poupiot, Sonia Albini, Daniel Stockholm, Laetitia Van Wittenberghe, Adeline Miranda, Nathalie Daniele, Inés Barthélèmy, Stephan Blot, Mai Thao Bui, Teresinha Evangelista, Isabelle Richard, David Israeli

**Author notes:** Corresponding author: David Israeli.

## Abstract

Duchenne Muscular Dystrophy (DMD), a muscle degenerative disease affecting young boys, arises from the loss of dystrophin. Current gene therapy approaches aim to restore a shortened form of dystrophin (microdystrophin) via Adeno-Associated Vector (AAV) delivery, but clinical studies show limited efficacy, emphasizing the need for improved strategies such as combined therapies. In this study, we identified lysosomal perturbations in the myofibers of DMD patients and animal models, an overlooked mechanism of cellular damage in muscular dystrophies. These were notably marked by the upregulation and recruitment of Galectin-3, a known biomarker of lysosomal membrane permeabilization, to damaged lysosomes, alongside alterations in lysosome number, morphology, and activation of endolysosomal damage response. Importantly, microdystrophin therapy in *Dmd^mdx^* mice fails to fully correct these damages. However, combining it with trehalose, a lysosome-protective disaccharide, significantly improves outcome, enhancing muscle function, histology and transcriptome. These findings highlight lysosomal damage as a novel mechanism in DMD pathogenesis and suggest that combining trehalose with gene therapy could enhance therapeutic efficacy.

**Teaser:** Lysosomal damage contributes to DMD pathology and is an interesting therapeutic target worth considering for current and future approaches.

## Introduction

Duchenne muscular dystrophy (DMD) is an X-linked recessive muscle disease due to the absence of dystrophin, the protein encoded by the DMD gene. It affects roughly 1/5000 boys and is the most common form of childhood muscular dystrophy. The loss of dystrophin in the skeletal myofiber leads to a destabilization of the dystrophin-associated glycoprotein complex, which connects the cytoskeleton to the extracellular matrix (Duan et al., 2021). This disruption triggers a cascade of pathological events including perturbations of calcium homeostasis, oxidative stress, mitochondrial dysfunction, chronic inflammation, fibro-adipocytic transformation, and consequently loss of muscle mechanical function (Allen et al., 2016). Current clinical management of DMD involves a multidisciplinary approach to address both muscular and extra-muscular manifestations of the disease (Roberts et al., 2023). Approved treatments include mainly glucocorticoids, which are recommended as early as possible (Zhang & Kong, 2021) and throughout the patient’s life (Bushby et al., 2010). Although glucocorticoids have shown a significant improvement in motor function, they remain merely a symptomatic treatment and can at best delay disease progression (McDonald et al., 2018). Besides, several therapeutic approaches under investigation aim at restoring a functional form of dystrophin. One of the most advanced and promising approaches is microdystrophin (µ-dystrophin) gene therapy, which involves delivering a shortened but functional variant of the dystrophin gene using a recombinant Adeno-Associated Vector (rAAV) (Chamberlain et al., 2023). This approach faces a key challenge, which is optimizing the therapeutic dosage. High doses may trigger severe immune responses and organs’ toxicity (Bradbury et al., 2023), while low doses may fail to achieve the desired therapeutic effect. Although one µ-dystrophin gene therapy (*delandistrogene moxeparvovec*) has recently gained FDA approval for ambulatory patients, functional improvements for patients have not replicated preclinical results (Mullard, 2023; Rind, 2024). The limited functional improvement observed in the clinical trials with µ-dystrophin could be attributed to several factors, primarily including the lack of certain functional domains found in full-length dystrophin (Mirouse, 2023). Therefore, µ-dystrophin gene transfer may not be able to reverse all the pathological mechanisms in the muscle. Identifying specific pathological mechanisms that remain uncorrected by µ-dystrophin gene therapy and exploring combinatorial approaches with complementary treatments targeting these mechanisms is crucial.

It is increasingly appreciated that the lack of dystrophin results not only in the mechanical instability of the skeletal myofiber but also in a vast spectrum of metabolic perturbations (Timpani et al., 2015). Among these, plasma dyslipidemia and perturbation of lipid metabolism were identified (White et al., 2020), whereas, in an investigation of expression of circulating microRNA in a large DMD cohort, we have recently showed perturbations of cholesterol metabolism in the dystrophic muscle in DMD (Amor et al., 2021; Israeli et al., 2022). Excessive cholesterol levels have frequently been associated with impaired lysosomal function, particularly in neurodegenerative diseases (Betuing et al., 2022; Udayar et al., 2022). We therefore hypothesized that cholesterol accumulation in the dystrophic muscle could be associated with impaired lysosomal function. To assess lysosomal function in DMD, we first evaluated the expression and localization of galectin-3 (Gal-3 or LGALS3), a known biomarker of lysosomal membrane permeabilization (LMP). We found that Gal-3 is significantly upregulated in the skeletal myofibers of DMD patients and various animal models, localizing to the lysosomes. LMP was also detected in a Limb-Girdle Muscular Dystrophy Type R5 (LGMDR5) mouse model and patient, suggesting a common perturbation across different types of muscular dystrophy. Using the *Dmd^mdx-4Cv^* mice, we detected and characterized various lysosomal perturbations in the dystrophic muscle, such as enlargement and increased number of lysosomes. Additionally, we observed activation of the endolysosomal damage response, which includes lysosomal replacement, repair and removal, and a perturbation of autophagy. Remarkably, LMP was exacerbated in dystrophic mice fed a cholesterol-rich diet and was not fully corrected by µ-dystrophin gene therapy. In contrast, gene therapy showed better corrected in the LGMDR5 mouse model when delivering the complete sequence of the missing gamma-sarcoglycan coding sequence, suggesting that the corresponding lysosomal dysfunction is reversible in other muscular dystrophies, but not by the sole µ-dystrophin in DMD. Subsequently, we selected trehalose, an FDA-approved compound known to restore lysosomal function, for use in combination with a suboptimal dose of AAV-µ-dystrophin gene therapy. The combined treatment resulted in improved correction of dystrophic parameters, including motor function, muscle histology, and transcriptome signature, compared to µ-dystrophin suboptimal gene therapy alone.

## Results

### Lysosome membrane permeabilization impairs lysosome function in muscle cells and can be detected with LGALS3 puncta assay

To investigate the hypothesis that cholesterol accumulation in the dystrophic muscle could be associated with impaired lysosomal function, we needed a method to quantify lysosomal damage. Several studies have recently focused on lysosomal dysfunction, in which LMP plays a central role (Lakpa et al., 2021). However, very few studies have investigated lysosome damage in muscle. One identified marker of LMP is gal-3 (or LGALS3), which is part of the galectins, a family of lectins that bind specifically to carbohydrates (Barondes et al., 1994). These lectins are present in the cytosol, nucleus, and extracellular matrix (ECM) and are translocated to the membrane of damaged lysosomes prior to their removal by the autophagy machinery (Aits et al., 2015). To analyze LGALS3 expression in a myogenic environment and its capacity to detect LMP, healthy human immortalized myoblasts were differentiated into elongated myotubes for 7 days, and then immune-stained for LGALS3. A diffuse expression of LGALS3 was observed in the cells (**Figure S1a**). Treatment with L-leucyl-L-leucine methyl ester (LLOMe), a lysomotropic agent that induces lysosome-specific membrane damage (Uchimoto et al., 1999), for 30 minutes at 2,5 mM, triggered the formation of LGALS3 positive puncta, a characteristic of damaged lysosomes. An overexpression of LGALS3 was also detected by western blot after 30 minutes to 1 hour of LLOMe treatment (**Figure S1b-c)**. Interestingly, LGALS3 puncta were drastically reduced 3 hours after treatment, and LGALS3 expression was restored to control level, demonstrating an efficient lysosomal repair by the cells. This pattern correlated inversely with the amount of LGALS3 released in the media, detected by enzyme-linked immunosorbent assay (ELISA) assay (**Figure S1d**), where a significant increase was detected in the media only after 3 hours of LLOMe treatment. We then aimed to check the effect of LMP on lysosomal function after LLOMe treatment. For that, we performed pH probing using a combination of two dextran conjugates (MW = 10 kDa), which are endocytosed by the cells, and accumulate selectively in the lysosomes. One dextran is conjugated with Oregon Green, whose emission is quenched under acidic environments such as the lysosomal lumen, and the other is conjugated with Tetramethylrhodamine (TMRM), whose emission in the red spectrum is pH independent (Fernandez-Mosquera et al., 2019). Monitoring the ratio between the green and red dextran-coupled fluorophores demonstrated a decrease of acidification after treatment with LLOMe for 30 minutes (higher green/red ratio), indicating an impact of LMP on lysosomal function in myotubes. Collectively, these data indicate that LGALS3 is expressed in muscle cells and is recruited specifically to damaged lysosomes. Immunostaining for LGALS33 could be therefore applied to detect LMP in skeletal muscle and to provide an insight into lysosomal function.

### LGALS3 is overexpressed in dystrophic muscle and correlates with lysosomal membrane permeabilization and lysosomal stress

After validating LGALS3 as a biomarker for LMP in muscle cells, we aimed at studying its expression and localization in DMD. First, Gal-3 expression at the mRNA and protein levels was evaluated in young 7-week-old *Dmd^mdx-4Cv^*mice. A significant upregulation was observed in both mRNA transcripts and protein in the gastrocnemius (GA) (mRNA: FC=32; protein: FC=14.4), tibialis anterior (TA) (mRNA: FC=21; protein: FC=22.6) and diaphragm (mRNA: FC=10; protein: FC=41.2) muscles of dystrophic mice compared to Wild-Type (WT) mice (**Figure 1a-c**). In addition, a significant increase of LGALS3 was also detected in the serum of *Dmd^mdx-4Cv^* mice compared to control mice (FC=1.8) (**Figure 1d**), suggesting a release of LGALS3 from the muscle. To evaluate whether LGALS3 is recruited to the damaged lysosomes in the context of LMP in the dystrophic muscle, cross-sections of TA muscles were immunostained for LGALS3 and the lysosome-associated membrane proteins 2 (LAMP2) which is commonly used as lysosome marker (Barral et al., 2022) (**Figure 1e**). Confocal immunofluorescence images showed a colocalization of LGALS3 puncta with LAMP2 inside the myofibers of young *Dmd^mdx-4Cv^*mice, indicating a substantial amount of LMP. However, LGALS3 and LAMP3-positive areas were also observed outside the myofibers, in regions consistent with inflammatory infiltrates characteristic of dystrophic muscle (**Figure 1e**). This aligns with prior reports attributing Gal-3 upregulation in DMD animal models primarily to LGALS3-positive infiltrating immune cells such as macrophage (Coulis et al., 2023; Marotta et al., 2009; van Putten et al., 2012).To confirm the inflammatory origin of these regions, co-immunostaining for LGALS3 and CD11b was performed on serial TA muscle cross sections (**Figure 1g**). An overlay of both staining (CD11b and LGALS3) was observed between the myofibers, confirming the presence of infiltrating macrophages, while LGALS3 staining was also evident within myofibers. This indicates that the upregulation of Gal-3 in muscle lysates reflects contributions from both immune infiltrates and myogenic sources. To specifically quantify LMP in myofibers, we measured LGALS3+LAMP2+ puncta within myofibers, excluding inflammatory areas using a segmentation method (**Figure S2a**). This analysis revealed a significant increase in LGALS3+LAMP2+ puncta in dystrophic muscle compared to WT controls (mean puncta = 9 for *Dmd^mdx-4Cv^* versus 1 in WT), confirming that LGALS3 overexpression in *Dmd^mdx-4Cv^* muscles is not solely due to macrophage infiltration but also reflects myogenic upregulation and recruitment to lysosome. These findings strongly suggest substantial LMP in dystrophic myofibers.

**Figure 1:**
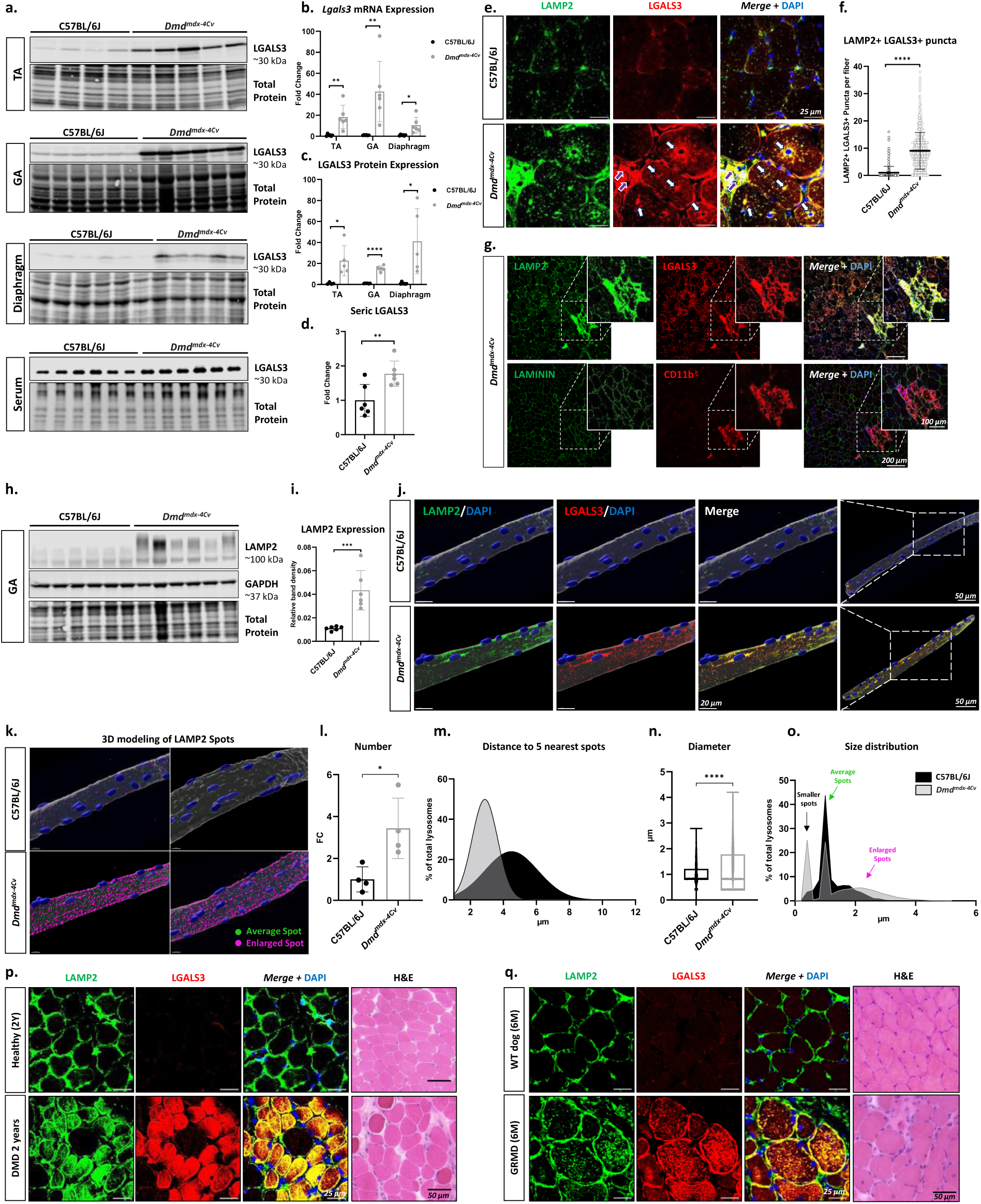
LGALS3 overexpression in dystrophic muscle correlates with LMP and lysosomal stress. **(a)** Western blot analysis of TA, GA, Diaphragm muscles and serum samples of WT and *Dmd^mdx-4Cv^* 7 weeks-old mice (n=4-6) for LGALS3. Total protein staining was used as loading control. (**b)** Relative Lgals3 mRNA expression in the TA, GA, and diaphragm, normalized to Rplp0. Scatter plots represent average ± SD of Fold change compared to WT mice (n=6). **(c)** Band intensity quantification of LGALS3 relative to total protein staining in TA, GA, and diaphragm muscles. Scatter plots show average ± SD of Fold change compared to WT mice (n=4-6). (**d)** Band intensity quantification of LGALS3 relative to total protein staining in serum. Data are represented as scatter plot of average ± SD of fold change relative to WT mice (n=6). (**e)** Representative confocal Images of TA muscle cross-sections immunostained for LAMP2 (green) and LGALS3 (red). White arrows indicate LAMP2+LGALS3+ puncta inside the myofibers, purple arrows indicate immune cells infiltration between the myofibers. (**f)** Quantification of double positive puncta (LAMP2+LGALS3+) in the myofibers of TA muscles of WT and Dmd^mdx-4Cv^ mice. Data are represented as dot plots with average ± SD (>400 myofibers from n=6). (**g)** Confocal images of serial transversal sections of TA muscles from *Dmd^mdx-4Cv^* mouse immunostained for LAMP2/LGALS3 and LAMININ/CD11b. Zoomed in images show inflammatory area of the dystrophic muscle. (**h)** Western blot analysis of GA muscle for LAMP2 (n=6). (**i)** Band density quantification of LAMP2 relative to GAPDH (n=6), represented as scatter plots (average ± SD). (**j)** Reconstructed 3D images of isolated myofibers from the flexor digitorum brevis (FDB) muscle of *Dmd^mdx-4Cv^*and WT mice, immunostained for LAMP2 and LGALS3. (**k)** 3D modeling of LAMP2 spots in single myofibers. Green spots show average sized lysosome (diameter <1.5 µm), and purple spots show enlarged lysosomes (diameter >1.5 µm). (**l)** Analysis of number of lysosomes per myofiber (n=4) (average ± SD of Fold-Change are represented). (**m)** Analysis of 5 nearest spots to each spot, represented as density plot (with non-linear fit gaussian curve applied). (**n)** Quantification of LAMP2 spots diameter (bow plots showing min to max data points). (**o)** Density plot showing distribution of LAMP2 spots diameter. (**p-q)** Representative confocal images of transversal sections from muscles of a DMD patient and a healthy control **(p)**, and a GRMD dog and a WT control **(q)**, immunostained for LAMP2 and LGALS3. Left panels show H&E labeling from the same muscle biopsies. An unpaired two-tailed t.test was used for statistical comparisons of WT and Dmd^mdx-4Cv^ groups. T.test p-value *p < 0.05, **p<0.01, ****p<0.0001.

After confirmation of LMP incidence in the dystrophic muscle, we sought to evaluate parameters of lysosomal stress. First, we evaluated lysosome number with a WB for LAMP2 in WT and *Dmd^mdx-4Cv^* muscles (**Figure 1h & S2b**). An overexpression of LAMP2 was observed in both GA (FC_GA_=3.9) (**Figure 1i**) and TA (FC_TA_=4.1) (**Figure S2b**). Moreover, a quantification of LAMP2 spots on transversal sections from TA muscles immunolabeled with LAMP2 showed a significant increase of spot number (mean=14.3 and 12.2 for *Dmd^mdx-4Cv^* and WT respectively) (**Figure S2c-d**), and spot size (FC=1.7) [**Figure S2e**), two main criteria of lysosomal stress. To further analyze lysosome morphology and number at the single myofiber level, Flexor Digitorum Brevis (FDB) muscle fibers were isolated from *Dmd^mdx-4Cv^* and WT mice, and immunostained for LAMP2 and LGALS3 (**Figure 1j**). Three Dimension re-construction of confocal stack images showed a significant accumulation of lysosomes in the *Dmd^mdx-4Cv^* myofibers, which co-localized with LGALS3, especially around the nuclei, indicating a substantial amount of LMP. However, not all lysosomes (LAMP+ spots) were positive for LGALS3, indicating the existence of a population of non-damaged lysosomes. A 3D modeling of LAMP2 spots (**Figure 1k**), showed a significant increase of lysosome number (FC_number_=3.4) (**Figure 1l**) and a denser distribution in the *Dmd^mdx-4Cv^* myofibers, as indicated by the distance to the 5 nearest spots (FC_5 nearest spots_=-1.65) (**Figure 1m**). An increase of the average spot diameter (FC_Diameter_=1.2) was observed in the *Dmd^mdx-4Cv^* myofibers (**Figure 1n**). When plotting the spots diameter on a density plot, three populations of LAMP2 spots were distinguished in the *Dmd^mdx-4Cv^*myofibers versus one main population in the WT myofibers (**Figure 1o**). Both WT and *Dmd^mdx-4Cv^* myofibers displayed a population of average size spots. *Dmd^mdx-4Cv^* myofibers showed additionally a population of enlarged spots, corresponding to typical damaged enlarged lysosomes, indicative of lysosomal stress and accumulation of undegraded materials. Interestingly, an additional population of smaller spots was detected in *Dmd^mdx-4Cv^*myofibers. These smaller spots may correspond to newly generated lysosomes and/or to proto-lysosomes, which are fragments of recycled lysosomal membranes following the elimination of damaged lysosomes (Y. Chen & Yu, 2017).

Collectively, these data indicate that LMP occurs in dystrophic muscle of *Dmd^mdx-4cv^* as detected by LGALS3+LAMP2+ puncta quantification. Furthermore, various signs of lysosomal stress, including lysosome enlargement and increased number and density, were detected in the myofibers of dystrophic mice, further supporting the hypothesis of lysosomal dysfunction in the dystrophic muscle.

### Lysosome membrane permeabilization persist with age, and is a common feature across different muscular dystrophies and models

To address whether LMP is only an early process in dystrophic pathogenesis since only young mice were evaluated above, or a persistent feature with disease evolution, 1 year-old *Dmd^mdx-4Cv^* and control mice were also studied. An upregulation of *Lgals3* transcripts (FC=29 for TA and 6.5 for GA) (**Figure S3a**), an overexpression of LGALS3 (FC=52 for TA and 23 for GA) (**Figure S3b-c**), and a high number of LGALS3+LAMP2+ puncta (**Figure S3d-g**), were detected in both TA and GA muscle (Mean =10.7 and 9.1 in *Dmd^mdx-^* versus 3.9 and 3.1 in WT, for TA and GA respectively). Interestingly, older WT mice displayed more LMP compared to young WT mice, indicating a possible degradation of lysosomal function overtime. Overexpression of LGALS3 in muscle fibers was also confirmed on biopsies of DMD patients of different ages (2 years and almost 4 years) (**Figure 1o & S3h**) and of 6-month-old *golden retriever muscular dystrophy* (GRMD) dogs (**Figure 1p**), with all biopsies showing a colocalization of LGALS3 and LAMP2 in the majority of myofibers of dystrophic muscles.

To evaluate if lysosomal degradation was a specific DMD feature, or whether it might be a common feature in muscular dystrophies, transversal GA muscle sections from a limb-girdle muscle dystrophy type R5 (LGMDR5 or gamma-Sarcoglycanopathy) mouse model (*Sgcg*^-/-^) were immunostained for LAMP2 and LGALS3 (**Figure S4a**). A significant increase of LAMP2+LGALS3+ puncta was observed in *Sgcg*^-/-^ muscle compared to WT control (mean = 8.3 vs 1.9 for control) (**Figure S4a-b**), indicating a substantial amount of lysosomal damage. Additionally, a muscle biopsy from an LGMDR5 patient was analyzed. A positive pattern of LAMP2+/LGALS3+ staining was detected inside the myofibers, although to a lesser extent than DMD muscle, possibly correlating the level of lysosomal damage and the severity of the dystrophic phenotype. Taken together, these findings show that LMP, which can be detected with LGALS3 recruitment to the lysosomes, might be a common cellular perturbation in muscular dystrophies.

### Activation of the endolysosomal damage response in *Dmd^mdx-4Cv^* muscles

Lysosomal stress, characterized by LMP and increase of lysosome size, is an activator of the endolysosomal damage response, which is coordinated, among other proteins, by LGALS3 (Jia et al., 2020). In physiological conditions, this response includes lysosomal replacement, repair and removal. We sought out to investigate whether and to which extent this response is activated in the dystrophic muscle.

First, we evaluated the lysosomal biogenesis pathway, mediated by the master transcription factor EB (TFEB). For that, we performed an immunostaining TFEB on muscle cross-sections of 7-week-old *Dmd^mdx-4Cv^* mice and WT controls (**Figure 2a**). Interestingly, a nuclear localization of TFEB was observed more abundantly in dystrophic mice compared to control (**Figure 2a-b**), indicating a translocation of TFEB into the nuclei and hence an activation of the lysosomal biogenesis machinery. Bulk RNA sequencing was performed on GA muscle of 7-weeks old WT and *Dmd^mdx-4Cv^* mice and analyzed to determine the expression of genes involved in lysosome function and structure. An upregulation of various TFEB target genes was observed in *Dmd^mdx-4Cv^* mice compared to WT controls, that include lysosomal enzymes, cathepsins, protons pumps and lysosomal membrane glycoproteins (**Figure 2c**). Among the cathepsins we further immunodetected the overexpression in the TA muscle of *Dmd^mdx-4Cv^* mice of cathepsins B and D (*CTSB* and *CTSD*) **(Figure S5a)**. While cathepsins overexpression in the dystrophic muscle had been previously shown, it remained mechanistically poorly understood (Kimura et al., 2024; Tjondrokoesoemo et al., 2016; Whitaker et al., 1983). Our data support that this upregulation may result from lysosomal stress activation of the TFEB pathway (Jia et al., 2020; Settembre et al., 2011). In addition to overexpression, in WT muscle cathepsins B and D exhibited a punctate pattern co-localizing with LAMP2, confirming their confinement within the lysosomes, while in dystrophic muscle, a higher expression of CTSB and CTSD was detected, with a more diffuse pattern, only partially co-localizing with LAMP2. This observation suggests a leakage of cathepsins from the lysosomal lumen, a hallmark of LMP, which can have a detrimental effect on cell homeostasis by promoting protein degradation, cellular damage, inflammation, and cell death (Xie et al., 2023).

**Figure 2.**
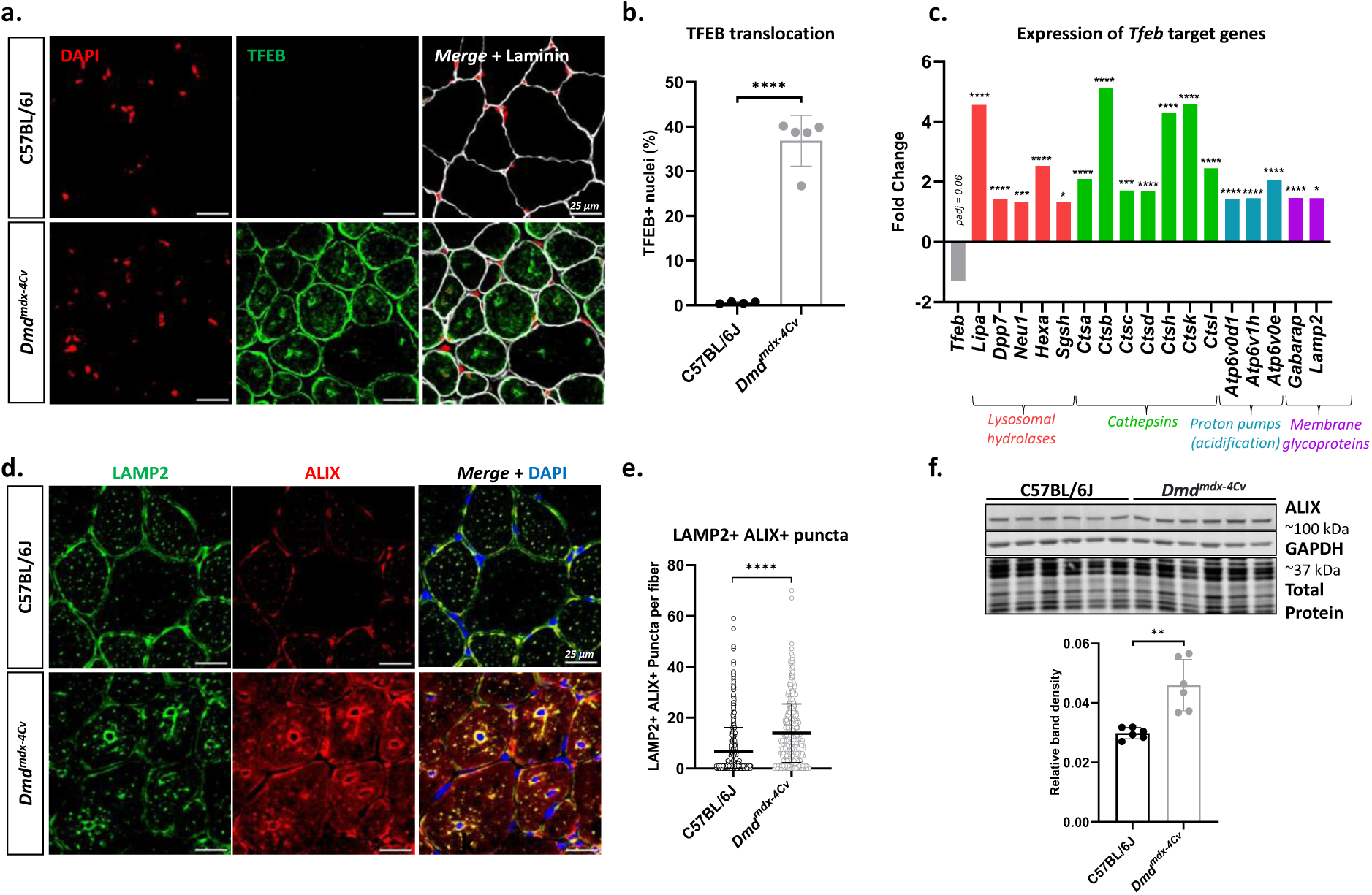
Activation of lysosomal biogenesis and repair pathways in *Dmd^mdx-4Cv^* mice. **(a)** Confocal images of serial transversal sections of GA muscle immunostained for TFEB and laminin. (**b)** Quantification of TFEB positive nuclei (n=5, average ± SD). (**c)** RNA sequencing analysis of GA muscle from 7-weeks old *Dmd^mdx-4Cv^* and control mice, displaying the expression of certain genes under the control of TFEB. Data are presented as fold change compared to WT controls (n=5). (**d)** Representative confocal images of TA transversal sections immunostained for ALIX and LAMP2. (**e)** Quantification of LAMP2+ ALIX+ positive puncta per myofiber. Data are presented scatter plots with average ± SD (>692 myofibers analyzed from n=6). (**f)** Western blot of TA muscle for ALIX, and band density quantification of ALIX relative to GAPDH (n=6), represented as scatter plots (average ± SD). An unpaired two-tailed t.test was used for statistical comparisons of WT and *Dmd^mdx-4Cv^* groups. T.test p-value *p < 0.05, ***p<0.001 ****p<0.0001.

Second, we evaluated the *Endosomal Sorting Complex Required for Transport* (ESCRT)-mediated lysosome repair, which is responsible for the quick repair of acute lysosomal leakages (Skowyra et al., 2018). Upon LMP, several subunits of the ESCRT complex are recruited to the lysosomal membrane in a Ca^2+^ dependent manner (Radulovic et al., 2018; Skowyra et al., 2018). Two of the recruited subunits are ALIX (ALG-2-Interacting Protein X or *Programmed Cell Death 6 Interacting Protein* - PDCD6IP) and the *Charged Multivesicular Body protein 4B* (CHMP4B). To evaluate this pathway, we performed an immunostaining of ALIX and LAMP2 on TA cross-sections (**Figure 2d**). A significant accumulation of ALIX puncta in *Dmd^mdx-4Cv^* mice, co-localizing with LAMP2, was detected (mean =13.9 and 6.8 for *Dmd^mdx-4Cv^* and WT respectively) (**Figure 2e**). An overexpression of ALIX protein was also detected in WB in TA muscles (**Figure 2f**). In addition, an increased colocalization of LAMP2 and CHMP4B was observed on TA muscle cross-sections compared to WT (**Figure S5c-d**), even though no overexpression of CHMP4B was detected at the protein level (**Figure S5e**). Together, these results suggest that lysosomal repair is active in the dystrophic muscle, through recruitment of the ESCRT complex.

Third, we aimed at evaluating lysophagy, which is the selective autophagy of damaged lysosomes where they are selectively captured by autophagosomes and subsequently degraded after fusion with intact lysosomes. As lysophagy is dependent on overall autophagic flux, we first aimed at assessing the general autophagy process. Whereas defects in autophagy have been previously reported in DMD, with a main role attributed to mTOR hyper-signaling (De Palma et al., 2012; Pal et al., 2014; You et al., 2024), no correlation with impaired lysosome function or structure was described. We performed a western blot on the TA for the *Microtubule Associated Protein 1 Light Chain 3 Beta* (MAP1LC3B or LC3B). An increase of LC3-I in the *Dmd^mdx-4Cv^* muscle (FC=1.7) was observed. However, the ratio between the lipidated form (LC3-II), generated during the autophagosome formation and representing active autophagy, and the non-lipidated LC3-I, was significantly reduced in *Dmd^mdx-4Cv^* muscles (FC=-1.9) (**Figure 3a**). This correlated with the increase of *Sequestosome 1* expression (SQSTM1 or p62) (**Figure 3b**), an autophagy substrate cargo protein known to be integrated into the autophagosomes and degraded during autophagy. These results confirmed a previous report in the mdx model (De Palma et al., 2012) and support the hypothesis of a decrease of the autophagic flux in the dystrophic muscle. To better understand the autophagy defects in dystrophic muscle, we performed an immunostaining of LAMP2 and LC3B on TA cross-sections (**Figure 3c**). A co-localization of LAMP2 and LC3B was notably detected around the nuclei in both WT and *Dmd^mdx-4Cv^* muscles, indicating lysosome-autophagosome fusion (Li et al., 2016; Yim & Mizushima, 2020). Analyses of LAMP2 and LC3B particles showed an increase of LAMP2+ (mean = 27 for *Dmd^mdx-4Cv^* vs 23.9 for WT) but not of LC3B+ particles (mean = 19.3 for *Dmd^mdx-4Cv^*vs 19.7 for WT, non-significant difference) in *Dmd^mdx-4Cv^* muscle compared to WT (**Figure 3d**). However, the number of LAMP2+LC3B+ particles, representing autolysosomes, was significantly decreased in *Dmd^mdx-4Cv^* muscle compared to WT (mean = 12.8 for *Dmd^mdx-4Cv^* vs 17.1 for WT). This analysis confirmed an increase of lysosomes (LAMP2+ LC3B-particles) ratio in the *Dmd^mdx-4Cv^* muscle compared to WT control (65% for *Dmd^mdx-4Cv^* vs 46% for WT) (**Figure 3e**), and a decrease of autolysosomes (LAMP2+ LC3B+ particles) ratio (35% for *Dmd^mdx-4Cv^* vs 54% for WT), which suggests a defect in lysosome-autophagosome fusion in the *Dmd^mdx-4Cv^* muscle. Moreover, to assess lysophagy specifically, we performed a co-immunostaining of SQSTM1 (labeling the autophagosomes (Pankiv et al., 2007)) and LGALS3 (labeling the damaged lysosomes) (**Figure 3f**). The increase of SQSTM1+ (mean= 37.6 for *Dmd^mdx-4Cv^*vs 22.6 for WT) and SQSTM1+LGALS3+ particles (mean= 21.1 for *Dmd^mdx-4Cv^* vs 7.8 for WT) in *Dmd^mdx-4Cv^* muscle indicated a significant activation of lysophagy (**Figure 3g**), and recruitment of the autophagic machinery to damaged lysosomes. Ratio analyses of total SQSTM1 particles (**Figure 3h**) confirmed an increase of lysophagy, represented by percentage of SQSTM1+LGALS3+ particles (55.2% for *Dmd^mdx-4Cv^*vs 32.8%), and a decrease of other autophagy-related pathways (percentage of SQSTM1+LGALS3-) in *Dmd^mdx-4Cv^* mice (44.8.% for *Dmd^mdx-4Cv^*vs 67.2%). Additionally, a co-immunostaining of LAMP2, LGALS3 and LC3B on TA cross-sections (**Figure 3i**) showed a colocalization of LAMP2 and LGALS3 with the autophagosome marker LC3B, indicating an ongoing lysophagy process, where damaged lysosomes are sequestered by the double membrane of the autophagosome, before fusing with remaining intact lysosomes (Maejima et al., 2013; Yang & Tan, 2023). Taken together, these data confirm an autophagy perturbation in the dystrophic muscle of *Dmd^mdx-4Cv^* mice, and notably a defect in lysosome-autophagosome fusion. Since, the fusion of autophagosome-lysosome requires intact acidic lysosomes, the substantial number of damaged lysosomes and the recruitment of the autophagic machinery to these damaged lysosomes in the context of lysophagy might be contributing to the autophagy blockage.

**Figure 3.**
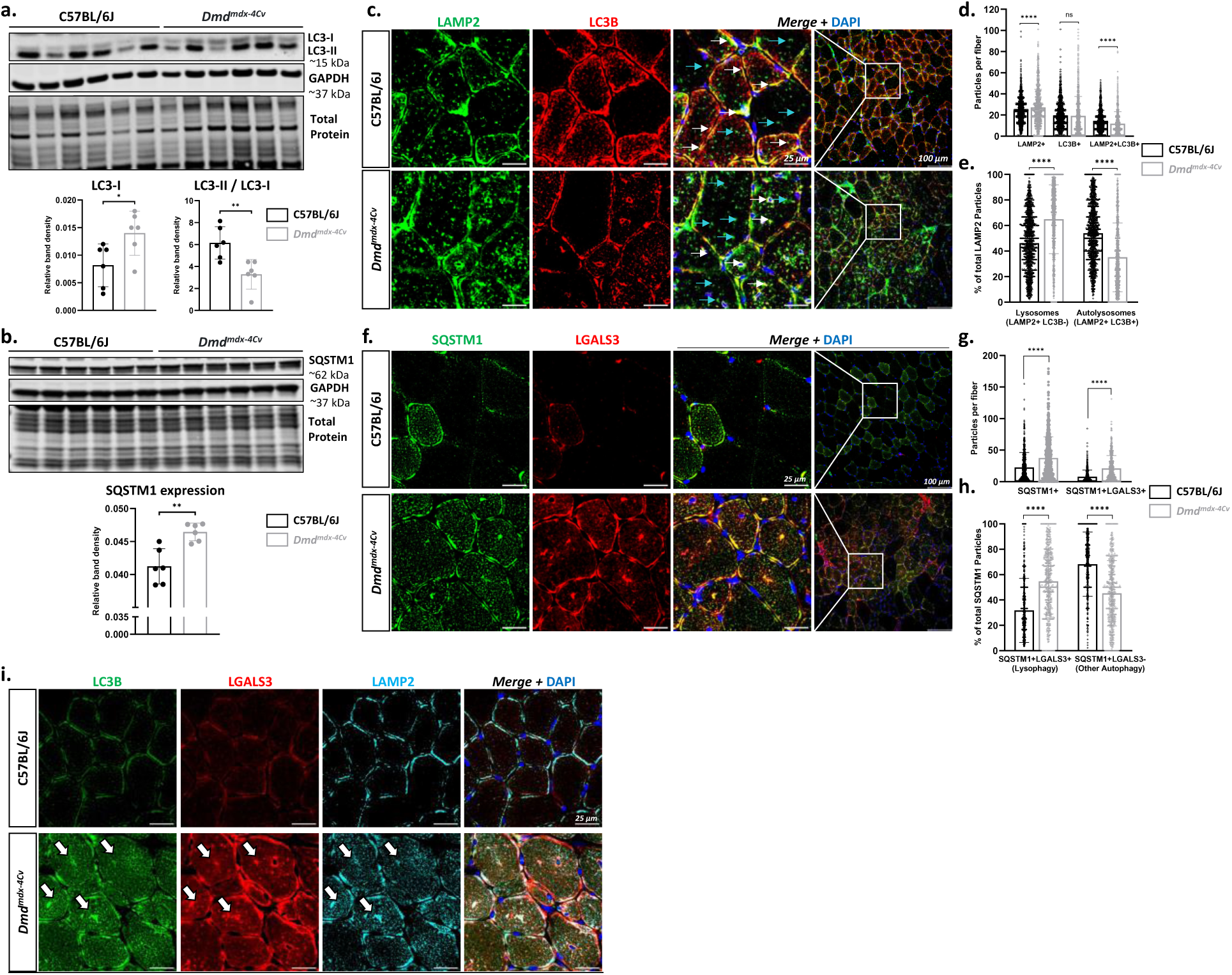
Autophagy and Lysophagy evaluation in *Dmd^mdx-4Cv^* muscle. (**a)** Western blot analysis of TA muscle for LC3, GAPDH and total protein staining are used as loading controls. Graphs show band density quantification of LC3-I and LC3-II/LC3-I ratio, normalized to GAPDH. Data are presented as scatter plots (average ± SD) (n=6). (**b)** Western blot analysis of TA muscle for SQSTM1, with GAPDH and total protein staining used as loading controls. Graph shows band density quantification of SQSTM1, normalized to GAPDH (average ± SD) (n=6). (**c)** Representative confocal images of TA transversal sections immunostained for LAMP2 and LC3B. White arrowheads depict LAMP2+LC3B+ particles representing autolysosomes, and blue arrowheads depict LAMP2+LC3B-particles representing lysosomes. **(d)** Scatter plots (average ± SD) showing quantification of LAMP2+, LC3B+ and LAMP2+LC3B+ particles in the myofibers (>1045 myofibers analyzed from n=6 per group). (**e)** Lysosomes and autolysosomes ratios were estimated by quantification of the percentages of LAMP2+LC3B-(lysosomes) and LAMP2+LC3B+ (autolysosomes) particles of total LAMP2+ particles per myofiber) (Scatter plots showing average ± SD). (**f)** Representative confocal images of TA transversal sections immunostained for SQSTM1 (p62) and LGALS3. (**g)** Scatter plots (average ± SD) showing quantification of SQSTM1+ and SQSTM1+LGALS3+ particles from immunostaining of TA muscle (>552 myofibers analyzed from n=6). (**h)** Ratios of active lysophagy and non-lysophagy related autophagy were estimated from the quantification of the percentages of SQSTM1+LGALS3+ (Lysophagy) and SQSTM1+LGALS3-(non-lysophagy autophagy) particles of total SQSTM1+ particles per myofiber (Scatter plots showing average ± SD). (**i)** Representative confocal images of TA transversal sections immunostained for LC3B, LGALS3 and LAMP2. White arrowheads depict LAMP2+LC3B+LGALS3+ representing damaged lysosomes sequestered by autophagosomes. An unpaired two-tailed t.test was used for statistical comparisons of WT and Dmd^mdx-4Cv^ groups. T.test ns=non-significant, p-value *p < 0.05, **p<0.01, ****p<0.0001.

### Lipid perturbations in *Dmd^mdx-4Cv^* mice correlate with damaged lysosomal system

We have previously shown perturbation of lipid metabolism, and especially accumulation of cholesterol in the dystrophic myofiber in DMD (Amor et al., 2021); but the consequences of these perturbations and their contribution to disease pathogenesis, including in relationship to lysosomal dysfunction, remained elusive. To further clarify the nature of these lipid perturbations, we evaluated muscle biopsies of young *Dmd^mdx-4Cv^*mice and a young DMD patient. Consistence with our previous study (Amor et al., 2021), significant accumulation of cholesterol was detected in the dystrophic muscle by filipin (free cholesterol) staining (**Figure S6a**). We additionally performed a lipidomic study in the GA muscle of young *Dmd^mdx-4Cv^* mice, which confirmed the cholesterol accumulation observed with filipin labelling (**Figure S6b-e**). Indeed, elevations of cholesterol, cholesterol esters and sphingomyelin were detected compared to WT mice (**Figure S6c-e**). Moreover, qPCR analysis in GA muscles of *Dmd^mdx-4Cv^* mice showed a dysregulation of major genes related to cholesterol homeostasis, such as HMG-CoA reductase (*Hmgcr*) involved in cholesterol synthesis, or the ATP-binding cassette transporter (*Abca1*) and apolipoprotein E (*ApoE*) involved in cholesterol efflux (**Figure S6f**). These data confirmed the perturbation of lipid metabolism and showed that cholesterol accumulation is an early process in DMD pathogenesis. To better elucidate the effect of cholesterol accumulation on the pathogenesis of DMD, and its potential link with lysosomal damage, young WT and *Dmd^mdx-4Cv^* mice were fed with either a cholesterol-rich high-fat diet or normal diet for 4 weeks (**Figure 4a**). An oil red O staining confirmed lipid accumulation in the liver of WT and *Dmd^mdx-4Cv^* mice under the high-cholesterol diet (HCD) (**Figure 4b**), but the HCD did not affect the body weight of the mice or the weight of muscles, liver, or heart (**Figure S7a-b**). Next, we evaluated the effect of the diet on the severity of the dystrophic phenotype. For that, muscle fibrosis in the diaphragm and the GA was analyzed by Sirius red labeling (**Figure 4c**). Quantification of Sirius red positive area showed a significant increase in fibrosis in *Dmd^mdx-4Cv^* mice on HCD compared to dystrophic mice on a standard diet (**Figure 4d-e**). However, no significant difference in fibrosis was observed between WT mice on HCD or standard diet in both the diaphragm and the GA (**Figure 4c-e**). This suggests that excess cholesterol in the diet may contribute to the exacerbation of the dystrophic phenotype. Nevertheless, no increase in muscle inflammation or sarcolemma permeabilization was caused by the HCD, as shown with CD11b and IgG labeling on GA cross-sections (**Figure S7c-e**). Additionally, we quantified the free cholesterol in the GA muscle (**Figure 4h**). *Dmd^mdx-4Cv^* mice on standard diet displayed higher muscle cholesterol accumulation than WT controls (FC_mdx-chow/WT-chow_=2.2), as expected. Interestingly, the cholesterol load increased in WT mice on HCD (FC_WT-HCD/WT-chow_=2), suggesting an accumulation of muscle cholesterol even in WT muscle. However, co-labeling of filipin (free cholesterol) and LAMP2 (lysosome) showed no significant localization of cholesterol in the lysosomes of WT muscle, contrary to *Dmd^mdx-4Cv^* muscle (**Figure 4f**), where a colocalization of filipin and LAMP2 was detected (**Figure i**). These results showed that HCD increases muscle cholesterol in both healthy and dystrophic muscles, but this accumulation was detected inside the lysosome only in the *Dmd^mdx-4Cv^* mice, suggesting an interplay between cholesterol accumulation and lysosomal dysfunction in dystrophic muscle. To further explore this hypothesis, we evaluated LMP following HCD with an immunostaining of LGALS3 and LAMP2 in the GA muscle of WT and *Dmd^mdx-4Cv^* (**Figure 4g**). An increase of LAMP2+LGALS3+ puncta was detected in *Dmd^mdx-4Cv^*mice on HCD compared to the group on standard diet (FC_mdx-HCD/mdx-chow_=1,8) (**Figure 4g&m**), which correlated with an upregulation of *Lgals3* (FC_mdx-HCD/mdx-chow_=1.75) and protein expression (FC_mdx-HCD/mdx-chow_=14.25) (**Figure 4j-l**). Interestingly, LAMP2+LGALS3+ spots were also detected in WT mice on HCD compared to the one on standard diet (mean = 3.2 and 0.1 for WT-HCD and WT-chow respectively), although these puncta were significantly less than those detected in the dystrophic muscle. Nonetheless, no significant increase of Gal-3 at the protein or mRNA level was observed in WT mice on HCD, suggesting that lysosomal damage may be minimal in WT mice on HCD. Taken together, these data indicate that excess cholesterol contributes to lysosomal damage in the muscle of both WT and dystrophic mice. However, the effect of the HCD on LMP is more significant in the dystrophic muscle, where lysosomal damage is already present and exacerbated further by cholesterol excess.

**Figure 4.**
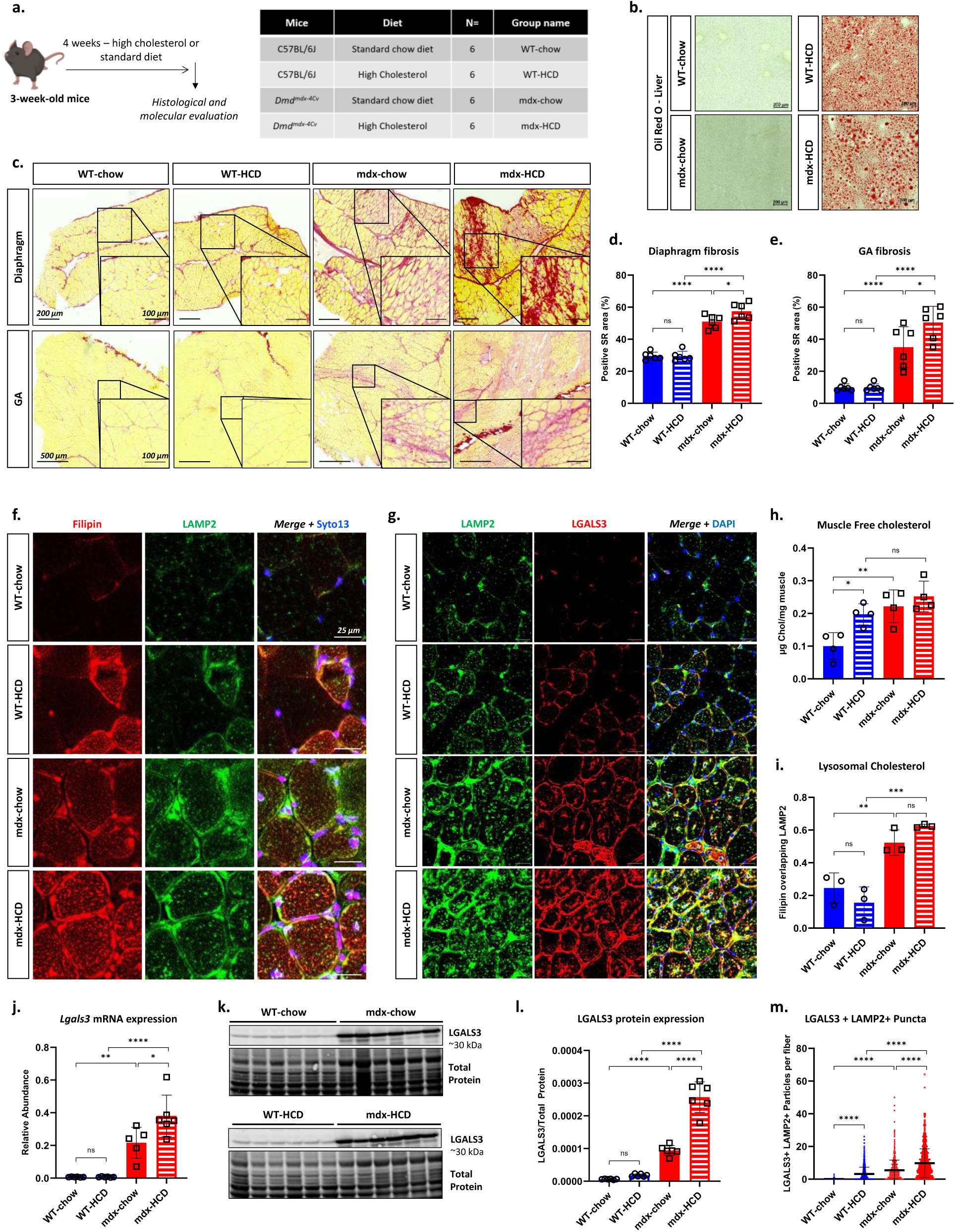
Lipid perturbations in *Dmd^mdx-4Cv^* mice correlate with LMP. (**a)** Schematic representation of study set-up. 3-week-old WT and *Dmd^mdx-4Cv^*mice were fed with a standard chow diet or a high cholesterol/high fat diet for 4 weeks. (**b)** Cross-sections of livers stained with Oil Red O, labeling non-polar lipids. (**c)** Sirius red staining of diaphragm and GA muscles cross-sections. (**d-e)** Quantification of fibrosis (Sirus red positive area) (n=6). (**f)** Representative confocal images of serial transversal sections of GA muscle labeled with filipin (Free cholesterol), Syto13 (nuclei) and immunostained with LAMP2 (green). (**g**) Representative confocal images of transversal sections of GA muscle immunostained for LAMP2 (green) and LGALS3 (red). (**h)** Scatter plot (average ± SD) displaying free cholesterol quantification in whole GA muscle lysates (n=4). (**i)** Co-localization analysis of Filipin signal overlapping LAMP2 signals, represented as scatter plot (average ± SD) of Manders’ Overlap coefficient (MOC) (n=3). (**j)** Relative Lgals3 mRNA expression in the GA normalized to Rplp0. Scatter plots represent average ± SD. (**k)** Western blot analysis of GA muscles (n=6) for LGALS3. Total protein staining is used as loading control. (**l)** Band intensity quantification of LGALS3 relative to total protein staining. Data are represented as scatter plot of average ± SD of fold change relative to WT mice (n=6). **m)** Quantification of double positive puncta (LAMP2+LGALS3+) in the myofibers of GA muscles. Data are represented as scatter plots with median (>784 myofibers analyzed from n=6). An ANOVA test with Tukey multiple comparison, or a Kruskal-Wallis with Dunn’s multiple comparisons (when values didn’t follow a normal distribution), were used for statistical comparisons. p-value *p<0.05, **p<0.05, ***p<0.001, ****p<0.0001, ns=non-significant.

### Correction of LMP following gene therapy varies between the modality of gene therapy approaches in dystrophic models

Following the identification of LMP in DMD, we set out to evaluate the effect of a current therapeutic approach to correct these damages, namely µ-dystrophin gene therapy. Six-week-old *Dmd^mdx^* mice were injected with an AAV9 viral vector encapsulating a µ-dys coding sequence under the control of a muscle promoter (spc5.12), and the analyses were conducted 6 weeks later. Two vector doses were used, a suboptimal dose (low) and a higher dose (high) known to be effective in this model (Bourg et al., 2022) (**Figure 5a**). A dose effect was noticed in terms of vector transduction assessed by quantification of viral copy number (VCN) (**Figure 5c**), transgene expression at the mRNA (**Figure 5d**) and protein levels (**Figure 5b&e**), with the high dose transducing nearly all GA muscle fibers (∼95% of positive myofibers) (**Figure 5b&e**). Additionally, a partial normalization of circulating biomarkers, creatine kinase (CK) and myomesin 3 fragments (MYOM3) was achieved, albeit not to WT levels (**Figure 5f-g**). A dose-dependent effect was observed in the reduction of fibrosis in the GA muscle, which was quantified from Sirius red labeling (**Figure 5b&i**). Still, even with the high dose of µ-dys, fibrotic area was not reduced to WT level in the GA muscle. Partial restoration of fibrosis correlated also with the rescue of global muscle force as evaluated with an escape test (**Figure 5h**). Indeed, the low dose did not increase muscle force compared to *Dmd^mdx^* controls, while the high dose resulted in a significant increase compared to non-treated mice, albeit not to WT level. Next, we investigated the correction of metabolic dysregulations. Labeling (**Figure 5b**) and quantification of free cholesterol (**Figure 5j**) were performed on GA muscles. An overcharge of free cholesterol was observed in *Dmd^mdx^*, as expected. Treatment with a low dose of AAV-microdystrophin resulted in a small but non-significant reduction in cholesterol. This reduction was greater with the high dose of AAV, where a non-significant difference from WT mice was obtained (**Figure 5j**). We then assessed the level of lysosomal damage following gene transfer. A 5.5-fold decrease of *Lgals3* mRNA expression in the GA was observed for mice treated with the high dose of µ-dys (FC_mdx-µ-dys/mdx-Phy-ser_=-5.5). A noticeable decrease of gal-3 at the protein level was also observed, but not to WT level. An immunostaining for LGALS3 and LAMP2 on GA cross-sections showed no reduction of LAMP2+LGALS+ puncta after treatment with low dose µ-dys, and only a subtle decrease after treatment with high dose µ-dys, compared with untreated mice (**Figure 5k&m**) (mean=7.8 for mdx versus 6.3 for mdx-µ-dys-high), indicating no or partial correction of LMP. These results suggests that treatment with a suboptimal dose of µ-dys is not sufficient to correct lysosomal damage, and a high dose of µ-dys can only reduce lysosomal damage but is still not sufficient for complete lysosomal correction. Following these results, we suspected that using µ-dys, rather than full length dystrophin, may result in incomplete lysosomal correction.

**Figure 5.**
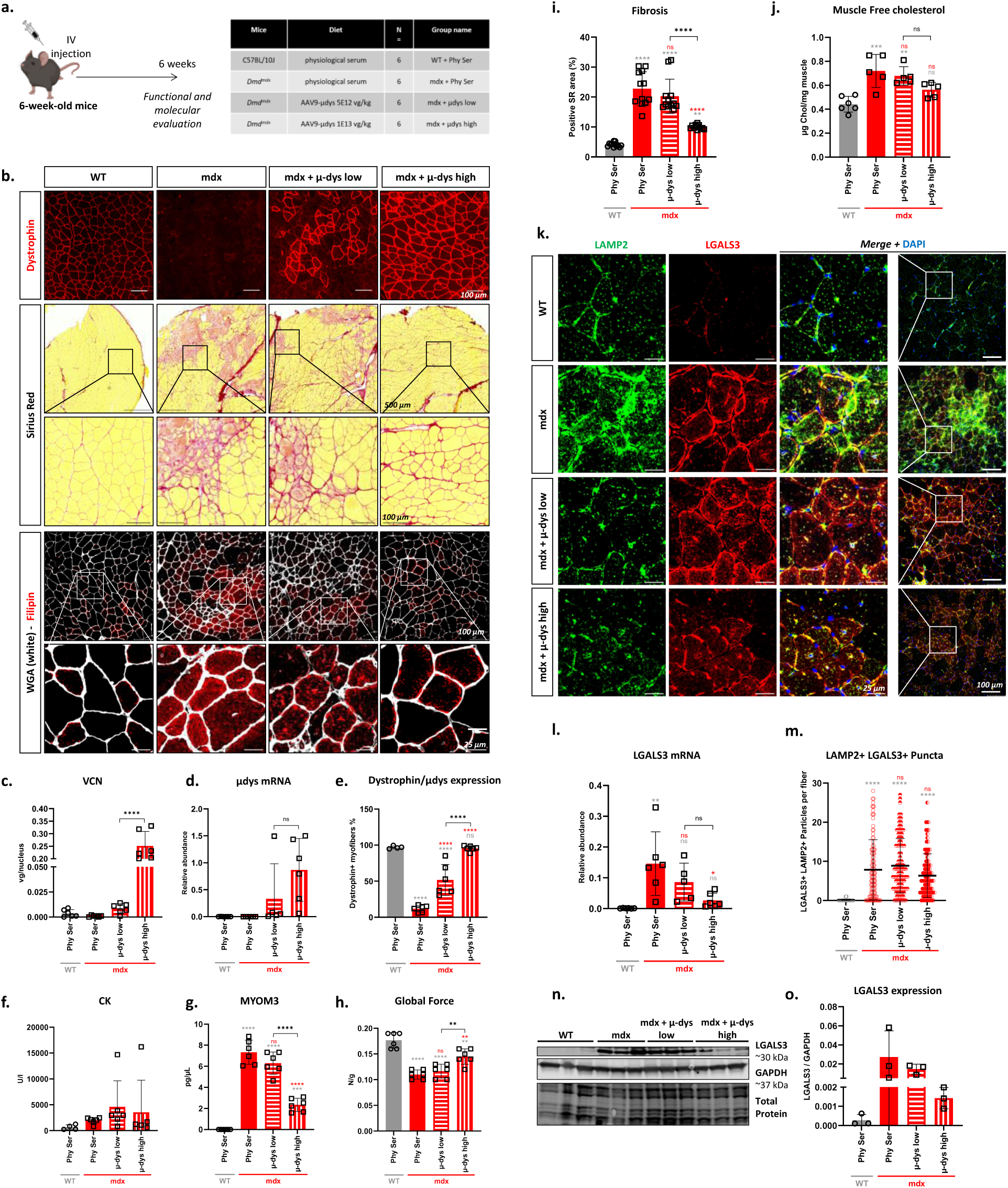
µ-dystrophin gene therapy does not entirely correct lysosomal damage in *Dmd^mdx^* muscle. (**a)** Schematic representation of study set-up. 6-week-old Dmd^mdx^ mice were inject intravenously with a rAAV encapsulating a µ-dystrophin encoding sequence under the control of spc512 promoter, at 2 doses of 5e12 vg/kg (low dose) and 1e13 vg/kg (high dose). (**b)** Histological characterization of GA muscle. Top panel: Dystrophin staining of GA cross-sections (DYSB antibody recognizes N-terminal part of dystrophin). Middle Panel: Sirus Red labeling of GA cross-sections. Bottom panel: Filipin (free cholesterol) and Wheat Germ Agglutinin (WGA) (labeling of nuclear and sarcolemma membranes) labeling of GA cross-sections. (**c)** Viral copy numbers (VCN) quantification in GA muscle. Data are presented as Scatter plot (average ± SD) of relative abundance to Rplp0. (**d)** Quantification of µ-dys transcripts in GA muscle. Data are presented as Scatter plot (average ± SD) of relative abundance to Rplp0. (**e)** Quantification of dystrophin positive fibers in GA muscle (The antibody used, DYSB, recognize the N-terminal of the µ-dys protein as well as the WT murine dystrophin). (**f)** Dosage of CK in serum of mice 6-weeks post-injection. Data are presented as scatter plot (average ± SD). (**g)** ELISA quantification of MYOM3 in the serum of the mice 6-weeks post injection. Data are presented scatter plot (average ± SD). (**h)** Global force evaluation by an escape test before sacrifice of mice. Data are presented as scatter plot (average ± SD) of global force normalized to mice body weight. (**i)** Quantification of fibrosis in the GA (Sirus red positive area) (n=6, two sections quantified per mouse). (**j)** Scatter plot (average ± SD) displaying free cholesterol quantification in whole GA muscle lysates (n=5). (**k)** Representative confocal images of transversal sections of GA muscle immunostained for LAMP2 (green) and LGALS3 (red). (**l)** Relative Lgals3 mRNA expression in the GA normalized to Rplp0. Scatter plots represent average ± SD (n=6). (**m)** Quantification of double positive puncta (LAMP2+LGALS3+) in the myofibers of GA muscles. Data are represented as scatter plots (average ± SD). (**n**) Western blot analysis of GA muscles (n=3) for LGALS3. GAPDH is used as loading control. (**o)** Band intensity quantification of LGALS3 relative to GAPDH. Data are represented as scatter plot of average ± SD (n=3). An ANOVA test with Tukey multiple comparison, or a Kruskal-Wallis with Dunn’s multiple comparisons (when values didn’t follow a normal distribution), were used for statistical comparisons. p-value *p<0.05, **p<0.05, ***p<0.001, ****p<0.0001, ns=non-significant. Grey Asterix represents relative comparison to ‘WT+Phy Ser’ group, red Asterix represent comparison to ‘mdx+Phy Ser’ group.

To investigate this hypothesis, we evaluated the effect of gene therapy on lysosomal correction in anothermodel for muscular dystrophy, notably LGMDR5, which in terms of pathophysiology is close to DMD. Of note, the gene therapy in that case is through the transfer of the entire gene and this investigation will help to define whether lysosomal dysfunction could be fully restored when using a complete copy of the missing gene. Using the *Sgcg^-/-^* KO mouse model, we delivered a complete copy of the human gamma-sarcoglycan gene and sought. *Sgcg^-/-^* KO mice were injected with a previously established optimal dose of 2e13 vg/kg (Israeli et al., 2019), which transduced nearly all muscle fibers with the human copy of the SGCG protein (**Figure S8a-e**). Histological, molecular, and functional evaluation showed the efficacy of the treatment, with correction of circulating biomarkers (**Figure S8f-g**) and muscle force (**Figure S7h**). Expression of *Lgals3* was also restored to WT levels after gene transfer. Number of LAMP2+LGALS3+ in the GA was also greatly decreased upon gene transfer (**Figure S7j-k**) (median= 5 for *Sgcg-/-* + PBS versus 1 for *Sgcg-/-* + AAV). Importantly, overall correction of dystrophic features and LMP was more effective in this model than µ-dys gene therapy in the *Dmd^mdx^* model.

Taken together, it appears that µ-dys gene therapy can only partially correct the muscle lysosomal damage, even when all myofibers are expressing µ-dys. This is of particular interest when compared to LGMDR5, where the transduction of all myofibers with the complete gamma-sarcoglycan protein is more effective at correcting LMP. It is therefore interesting to target specifically these damages in DMD, in combination with µ-dys gene therapy, in order to achieve lysosomal correction and improve overall reversion of dystrophy, as seen in gene therapy for *Sgcg^-/-^* KO mice.

### Trehalose-mediated lysosomal correction improves µ-dystrophin gene therapy at a suboptimal dose in a combined therapy setup

After the identification of lysosomal damage in the dystrophic muscle, and the insufficiency of the µ-dys gene therapy alone to correct these damages, we designed a combined therapy setup where a suboptimal dose of rAAV-µ-dys would be administrated with a drug capable of alleviating lysosomal dysregulations. For that, trehalose, an FDA-approved dietary disaccharide was selected, as it has been shown to activate lysosomal biogenesis and alleviate lysosomal damage in various neurodegenerative diseases (Palmieri et al., 2017; Rusmini et al., 2019). Trehalose was administered alone (2% in drinking water) and in combination with a suboptimal dose of rAAV-µ-dys in *Dmd^mdx-4Cv^* mice. To this end, five groups of male mice were included in this study (**Figure 6a**). Trehalose was diluted in the drinking water and administered from three weeks of age until the end of the study four weeks later. Recombinant AAV-µ-dys was injected intravenously (IV) one week after the beginning of treatment with trehalose. First, we assessed whether trehalose treatment affected muscle transduction or transgene expression. Quantification of VCN and mRNA transcripts in the GA and TA muscles showed no significant differences between the 2 groups injected with the vector with or without trehalose (**Figure S9a-b**). No significant difference was noticed in the number of dystrophin positive fibers as well (**Figure S9c-d**). Serum was collected at the end of the study before functional evaluation, and different circulating biomarkers were quantified. CK levels were decreased significantly compared to control *Dmd^mdx-4Cv^* reaching non-significant difference from WT mice only for mice treated with the combined therapy (FC_mdx-µ-dys/mdx_=-2.7, FC_mdx-µ-dys+TR/mdx_=-5.5) (**Figure 6c**). MYOM3 levels were significantly decreased with Trehalose treatment alone (FC_mdx-TR/mdx_=-1.3), but still not normalized to WT level even with the combined therapy (**Figure 6d**). As for the global muscle strength assessed by an escape test, a slight improvement in muscle strength compared to *Dmd^mdx-4Cv^*was observed in mice treated with trehalose alone or AAV-microdystrophin alone (**Figure 7e**), albeit not significantly (**Figure 6e**). Importantly, only mice treated with the combined therapy showed a significant improvement in muscle strength compared to the untreated *Dmd^mdx-4Cv^* mice and showed no longer a significantly different strength from the WT group (**Figure 6e**). We then performed a histological characterization, following the different treatments (**Figure 6b**). H&E staining of GA transversal section showed improvement of muscle histology for both trehalose and µ-dys only treated mice compared to non-treated *Dmd^mdx-4Cv^* mice (**Figure 6b**). However, a better correction was achieved in the combined therapy group, where myofiber size distribution was best restored (**Figure 6f**), and the centronucleation index decreased (**Figure 6g**). Fibrosis was assessed by Sirius red labeling in the diaphragm and GA, which are among the most fibrotic muscles in this mouse model (**Figure 6b**). As for fibrosis in the GA, the trehalose treated mice showed a significant reduction, and the combined therapy group had a significantly lower fibrosis than the µ-dys only treated group (**Figure 6b&h**). In the diaphragm, neither Trehalose nor µ-dys treatments alone decreased fibrosis (**Figure 6b&i**). Only the combination of treatments reduced fibrosis significantly in this muscle. In addition, myofiber necrosis was evaluated in the GA by mouse IgG labeling (**Figure 6b&j**). A significant reduction of IgG positive myofibers was observed in both trehalose and µ-dys treated mice, with a better correction in trehalose treated mice versus µ-dys-only (non-significant difference from WT controls). The best correction was attained in the combined therapy group, where very few IgG positive myofibers were detected, indicating a better integrity of myofibers sarcolemma. CD11b staining was also performed to assess the inflammation level in the GA muscle (**Figure 6b**). A significant decrease of inflammatory areas was observed with both trehalose and µ-dys treatments, with the combined therapy reaching the best correction (most significant decrease compared to non-treated *Dmd^mdx-4Cv^*).

**Figure 6.**
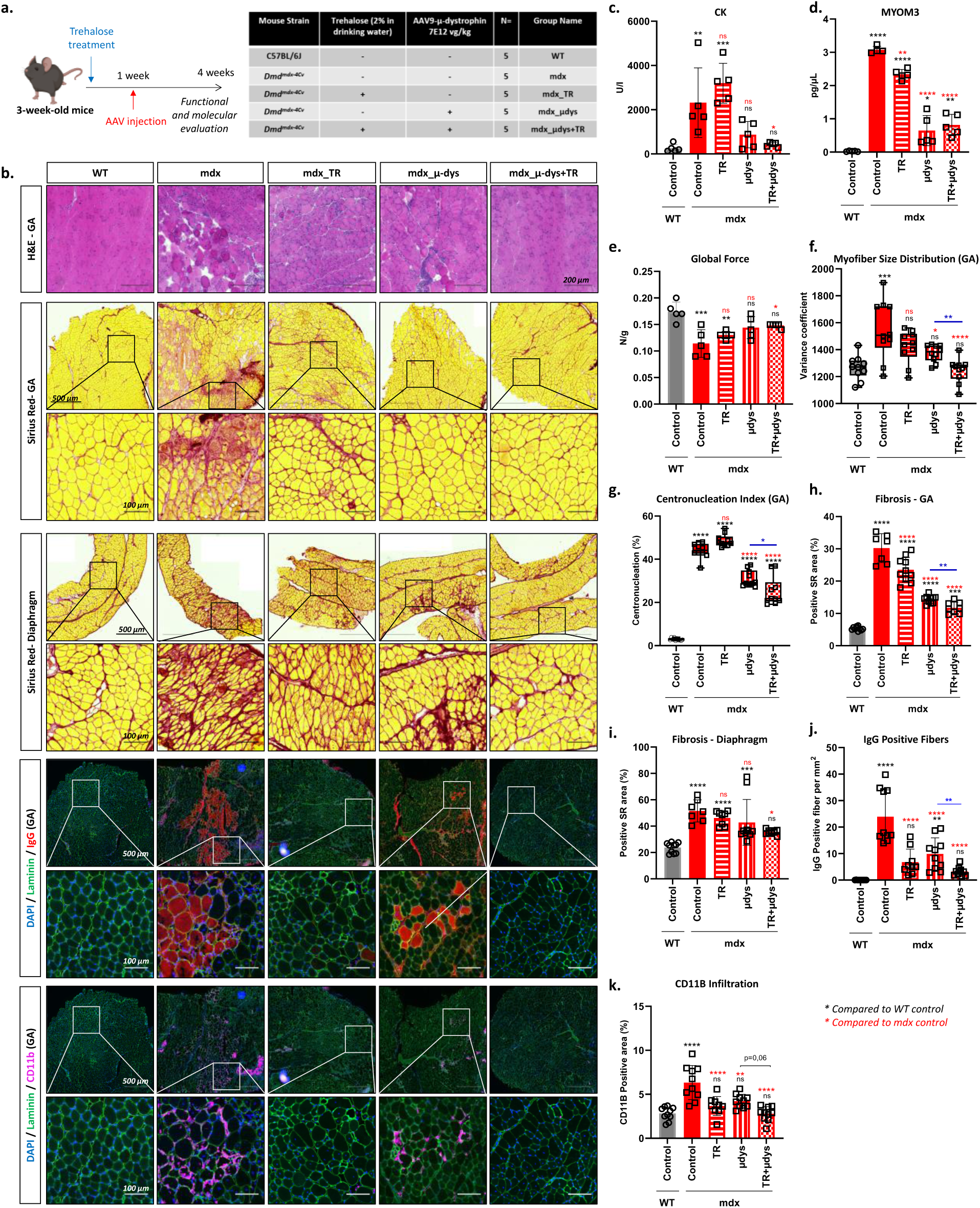
Trehalose treatment restore some dystrophic parameters and improve µ-dystrophin gene therapy in *Dmd^mdx-4Cv^* mice. (**a**) Schematic representation of study set-up. *Dmd^mdx-4Cv^* mice were started on trehalose treatment (2% dilution in drinking water) at 3-weeks. For Group 4 and 5, mice were injected intravenously with a rAAV9 encapsulating the µ-dys sequence at a 7e12 vg/kg dose. (**b**) Histological characterization of muscles. Top panel: H&E labeling of GA muscle cross-sections. Middle Panel: Sirius red labeling of diaphragm and GA muscle cross-sections. Bottom Panel: Representative images of GA muscle serial cross sections immunostained for mouse immunoglobulin (IgG), laminin and CD11b. (**c**) Dosage of CK in serum of mice before escape test. Data are presented as scatter plot (average ± SD, n=5). (**d**) ELISA quantification of MYOM3 in the serum of the mice before escape test. Data are presented scatter plot (average ± SD, n=5). (**e**) Global force evaluation by an escape test before sacrifice of mice. Data are presented as scatter plot (average ± SD, n=5) of global force normalized to mice body weight (N/g). (**f**) Analysis of myofiber size distribution in the GA muscle. Data presented as box plots (Median + minimum/maximum) show variance coefficient calculated as [standard deviation of the muscle fiber size/mean of muscle fiber size] *1000. (g) Centronucleation Index in the GA muscle. Data are presented as box plots (Median + minimum/maximum). h-i) Fibrosis analysis of GA and diaphragm, done by quantification of Sirius red positive area on transversal sections. Data are presented as scatter plots (average ± SD). (**j**) Quantification of permeabilized fibers positive for mouse IgG staining. Data are presented as scatter plot (average ± SD) of number of positive fibers normalized to muscle cross-sectional area. (**k**) Evaluation of CD11b+ cells infiltration in the muscle, done by quantifying CD11b area normalized to muscle cross-sectional area. Data are presented as a scatter plot (average ± SD). An ANOVA test with Tukey multiple comparison was used for statistical comparisons (compared to WT control and mdx control). ANOVA p-value *p<0.05, **p<0.05, ***p<0.001, ****p<0.0001, ns=non-significant. Black Asterix represents relative comparison to ‘WT control’ group, red Asterix represent comparison to ‘mdx control’ group. An unpaired two-tailed t.test was used for statistical comparisons between mdx-µ-dys and mdx_µ-dys+TR groups. T.test p-value *p < 0.05, **p<0.01.

**Figure 7.**
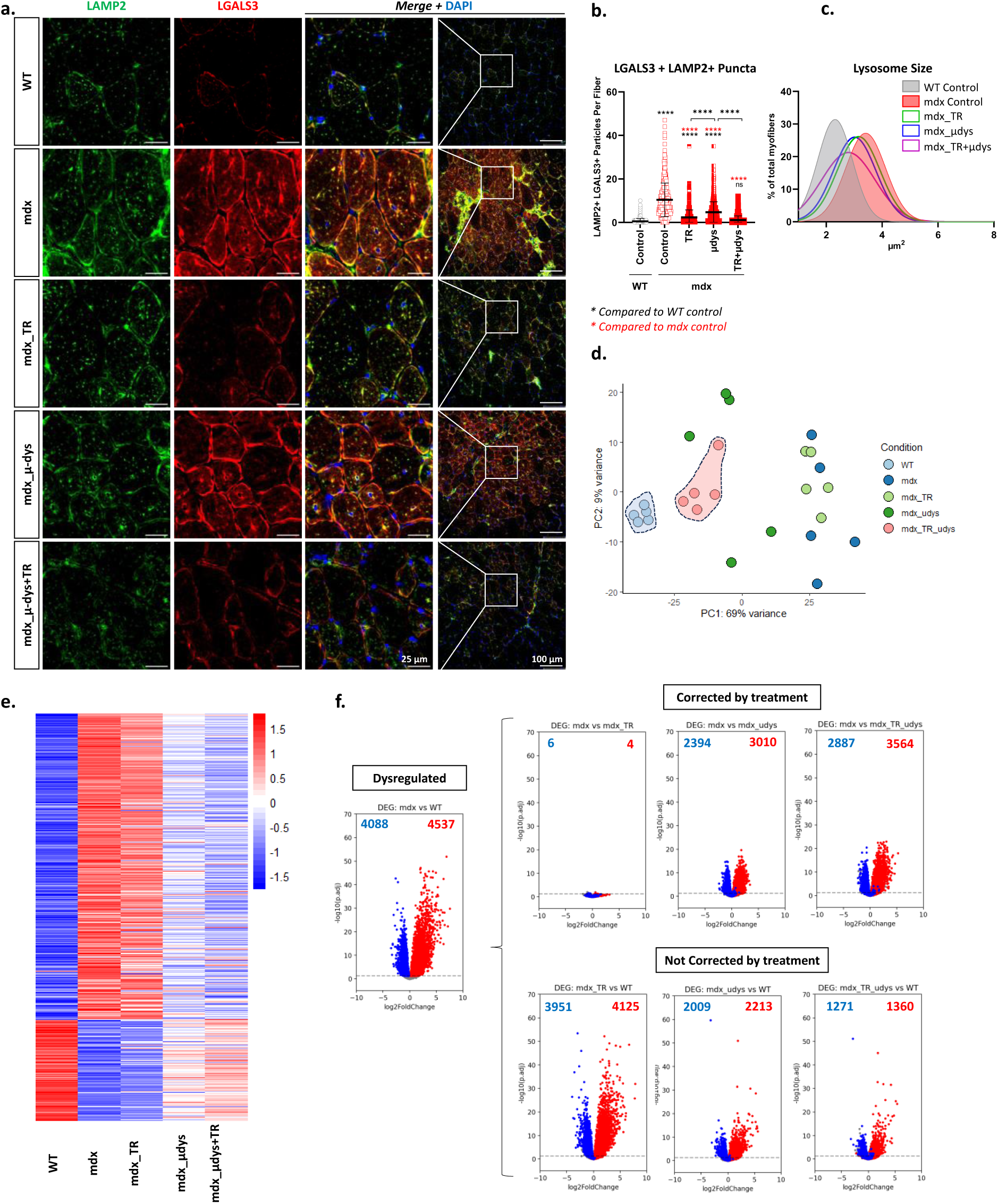
Trehalose treatment corrects lysosomal damage and improve transcriptome correction with µ-dys. (**a**) Representative confocal Images of transversal sections of TA muscle immunostained for LAMP2 (green) and LGALS3 (red). (**b**) Quantification of double positive puncta (LAMP2+LGALS3+) in the myofibers of TA muscle. Data are represented as scatter plots with (average ± SD) (>538 myofibers analyzed from n=5). (**c**) Density plot with non-linear fit gaussian curve applied, showing distribution of average spot size per myofiber (>538 myofibers analyzed from n=5). (**d-f**) Comparison of global transcriptomic changes in GA muscle. (**d**) Principal Component Analysis (PCA) plot using the expression level of 10000 genes with highest variance. Gene expression was first normalized by size factors and transformed by variance stabilizing transformation using the DESeq2 RStudio package. (**e**) Heatmap presenting the log2 fold change (log2FC) in comparison to WT muscle for all 8625 DEG found in mdx muscle (compared to WT). The log2FC values are illustrated in row Z-scores, colored from blue to red, arranged from lowest to highest. **(f**) Volcano plots of multiple comparisons illustrating transcriptomic changes before and after different treatments. As a reference, 4088 downregulated and 4537 upregulated DEGs found in mdx control were colored blue and red, respectively, in all volcano plots. Among these DEGs, the number of DEGs found to be significantly different in each pair-wise comparison were labeled in the upper corners. Right upper panel shows volcano plots comparing mdx control to treated groups, in which DEGs are correctly restored after treatment. Right lower panel shows treated groups compared to WT control, in which significant DEGs are the genes that are not or incompletely restored after treatment. An ANOVA test with Tukey multiple comparison was used for statistical comparisons in (**b**) and (**c**). ANOVA p-value ****p<0.0001. Black Asterix represents relative comparison to ‘WT control’ group, red Asterix represent comparison to ‘mdx control’ group.

Following the demonstration of better outcomes with the combined treatment, we wanted to assess the effect of trehalose on lysosomal damage and dysregulated molecular pathways. To this end, we immunostained LAMP2 and LGALS3 on TA muscle transversal sections (**Figure 7a**). A clear increase in LAMP2+LGALS3+ puncta was detected in *Dmd^mdx-4Cv^* muscle (mean = 10.4 and 0.5 for mdx and WT respectively), (**Figure 7b**). These puncta were decreased with µ-dys treatment (mean = 4.7) but decreased more significantly with the trehalose treatment (mean = 2.3). A correction to WT level was only achieved for the group with the combined therapy (mean = 1.1) (**Figure 7b**). Analyses of lysosomal enlargement were also done from LAMP2 immunostaining (**Figure 7c**). A clear enlargement was observed in *Dmd^mdx-4Cv^* muscle compared to WT, as seen on the density plot. Lysosomal size was significantly reduced with µ-dys or trehalose treatments. Nevertheless, the best correction was also best obtained in the mice treated by the combined therapy.

To assess the effects of different treatments on the reversion of transcriptomic perturbations in the dystrophic muscle, we performed analysis of bulk RNA-sequencing on the GA muscles. On the first two principal components (PCs) of the PCA (**Figure 7d**), the group treated with trehalose clustered around the untreated *Dmd^mdx-4Cv^* group, and the group treated only with µ-dys were widely distributed on the PCA. Importantly, only the group treated with µ-dys+Trehalose was tightly clustered close to the WT group. Next, we investigated the evolution of all differentially expressed genes found in *Dmd^mdx-4Cv^* untreated group (DEGs, comparison: untreated *Dmd^mdx-4Cv^* versus WT, padj<0.05) upon different treatments (**Figure 7e-f**). Similar to PCA, heatmap of the total 8625 DEGs showed comparable profiles between the trehalose treated and untreated *Dmd^mdx-4Cv^*group while µ-dys-alone treatment shifted the heatmap signature closer to WT (**Figure 7e**). Strikingly, the closest profile to WT was achieved with the combined therapy. Among the 4088 downregulated and 4537 upregulated DEGs identified in *Dmd^mdx-4Cv^* muscle, only 6 (0.15%) upregulated and 4 (0.08%) downregulated DEGs were restored by trehalose, while µ-dys alone restored 2394 (58.5%) upregulated and 3010 (66.3%) downregulated DEGs. Importantly, the combination of trehalose and µ-dys restored 2887 (70%) downregulated and 3564 (78.5%) upregulated DEGs. In addition, expression of more genes was partially or completely normalized by the treatment of µ-dys+trehalose compared to µ-dys alone. A total number of 2009 downregulated and 2213 upregulated DEGs were incompletely restored with µ-dys, compared to 1271 downregulated and 1360 upregulated DEGs for µ-dys+trehalose. We also performed gene set enrichment analysis (GSEA) to evaluate the correction of known dysregulated pathways in the dystrophic muscle, such as inflammatory response, calcium signaling, nitrogen species metabolic process and apoptosis (**Figure S10a-b**). Metabolic pathways like cholesterol homeostasis, lipid storage, endocytosis and MTORC1 signaling (**Figure S10c**) were also evaluated. Strikingly, these pathways were all significantly closer to normal with the combined treatment of µ-dys+Trehalose compared to µ-dys alone. Taken together, these data suggest that combining trehalose and µ-dys treatments significantly improve pathophysiology, lysosomal damage and transcriptomic dysregulation compared to suboptimal dosed gene therapy standalone treatment.

## Discussion

The prevailing dogma of DMD pathophysiology emphasizes the mechanical destabilization of the skeletal myofiber (Bez Batti Angulski et al., 2023; Duan et al., 2021). Accordingly, in the absence of dystrophin, skeletal myofibers are prone to contraction-induced damage, leading to sarcolemma disruption. Recently, however, there has been growing interest in the metabolic disturbances associated with DMD (Timpani et al., 2015), particularly mitochondrial dysfunction (Casati et al., 2024; Sanson et al., 2020; Timpani et al., 2015; Vu Hong et al., 2022). Additionally, several studies noted lysosomal perturbations in DMD (Gelman et al., 1981; Kominami et al., 1987; Pal et al., 2014; Sano et al., 1988), although the nature and consequences of these disturbances remained poorly understood. Interestingly, perturbations of lipid metabolism were also reported (Sun et al., 2023). Particularly, we and other have recently detected perturbations of cholesterol metabolism (Amor et al., 2021; Srivastava et al., 2017; White et al., 2021). Such disruptions of cholesterol metabolism are often linked with lysosomal damage, in both neurodegenerative and lysosomal storage diseases (Betuing et al., 2022; Kacher et al., 2022; Schulze & Sandhoff, 2011; Udayar et al., 2022; Whyte et al., 2017). The lysosomal damage biomarker LGALS3 (Aits et al., 2015) has been found to be upregulated in the dystrophic muscle in the *Dmd^mdx^* mouse model (Marotta et al., 2009). This upregulation was initially attributed to invading inflammatory mononucleated cells (van Putten et al., 2012), because Gal-3 is highly expressed in inflammatory macrophages (Coulis et al., 2023). However, recent studies suggest the expression of Gal-3 in muscle fibers and its requirement for efficient muscle regeneration (Cerri et al., 2021). We therefore set in the present study to investigate the expression of LGALS3 in the dystrophic muscle and confirmed not only its upregulation but also its localization to the enlarged lysosome of dystrophic myofibers. Importantly, the sole evaluation of Gal-3 expression at the mRNA or protein levels is not sufficient to estimate lysosomal damage, as infiltrating macrophages (Gal-3 positive) are the majority of immune cell infiltrate in dystrophic muscles. Therefore, it is important to quantify specifically LGALS3+LAMP2+ puncta inside the myofibers, to have a reliable estimation of lysosomal contribution to pathology and correction with various treatments.

We further validated and characterized lysosomal perturbations in dystrophic muscle, revealing increased number and size of lysosomes, activation of endolysosomal damage response, including replacement, repair and removal of damaged lysosomes. These processes are coordinated not only by the upregulation of LGALS3 (Jia et al., 2020) but also by the nuclear translocation of the TFEB transcription factor (Settembre et al., 2011) and the upregulation of its lysosomal target genes, such as the cathepsins, which happen to be upregulated in the dystrophic muscle (Kimura et al., 2024; Sano et al., 1988; Tjondrokoesoemo et al., 2016; Whitaker et al., 1983). Additionally, we identified autophagy defects in the dystrophic muscle. Although previous studies have already reported autophagy defects in DMD, these were mainly attributed to mTOR hyperactivation(De Palma et al., 2012; Pal et al., 2014; You et al., 2024). We found reduced rate of autophagosome-lysosome fusion, as well as increased lysophagy. Of note, lysosomal perturbations in skeletal muscle have been more widely studied before in the context of glycogen storage type II (GSDII/Pompe) disease. Interestingly, GSDII muscle is characterized by a blockage of autophagic flux with an impaired autophagosome-lysosome fusion (Do et al., 2024), despite a drastic downregulation of mTOR signaling. This autophagic blockade in GSDII can be reversed by the overactivation of TFEB (Spampanato et al., 2013). This suggests that also in DMD autophagic perturbation may result not only from mTOR upregulation but also be secondary to lysosomal dysfunction due to reduced autophagosome-lysosome fusion. Collectively, our data support a model where autophagic perturbation in the dystrophic DMD muscle results from a decreased availability of functional lysosomes, which should serve for autophagosome-lysosome fusion. This deficiency is partially mitigated by TFEB-induced lysosomal biogenesis. Additionally, we showed that lysophagy, aiming to remove compromised lysosomes, is highly activated in the dystrophic muscle. The upregulation of one selective form of autophagy, in this case lysophagy, could be limiting the capacity of other selective autophagy (Germain et al., 2024) .

To directly investigate the link between cholesterol accumulation and lysosomal damage, mice were fed a high-cholesterol diet (HCD). While increased cholesterol load in the skeletal muscle of HCD-fed mice were observed in both WT and dystrophic mice, abnormal cholesterol accumulation, lysosome damage and fibrosis were significantly more pronounced in the muscle of the HCD-fed dystrophic mice compared to those fed a standard diet, and HCD-fed healthy control mice. This indicates a direct causative relationship in this DMD model between elevated cholesterol, lysosomal damage and the exacerbation of the dystrophic parameters.

We then set out to test the effect of gene therapy on muscle lysosomal damage. As expected, a dose response relationship was observed, with LMP, evaluated by the LGALS3+LAMP2+ puncta, decreasing as the injected dose of AAV µ-dys increased. However, significant residual lysosomal damage persisted even in mice treated with a high dose of AAV µ-dys, which transduces nearly 100 % of the myofibers. Strikingly, similar gene therapy treatment of the LGMDR5 model, with the full-length gamma sarcoglycan transgene (Israeli et al., 2019) resulted in nearly complete reversion of LMP, indicating highly efficient lysosomal correction by gene therapy in this model. Trehalose was selected for the combined therapy setup due to its good safety profile and affordability as a dietary disaccharide (A. Chen & Gibney, 2023), and more importantly its previous testing in clinical trials for neurodegenerative disease (Pupyshev et al., 2022) and oculopharyngeal muscular dystrophy (OPMD) (Argov et al., 2015; Davies et al., 2006). Additionally, recent studies have shown that trehalose promotes lysosomal biogenesis and autophagy (Rusmini et al., 2019). Our experimental design enabled a comparison between the administration of trehalose alone, a suboptimal dose of AAV-µ-dys alone, and their combination. Notably, trehalose alone demonstrated significant benefits, particularly in reversing LMP (LAMP2+LGALS3+ puncta) and reducing myofiber permeabilization and necrosis (IgG staining). This highlights the positive effect of trehalose on lysosomal dysfunction and suggests a direct association between lysosomal damage and myofiber necrosis. Importantly, the combination of µ-dys gene therapy and trehalose yielded the best therapeutic results. In the GA muscle of treated mice, we observed correction of myofibers’ size distribution, improvement in the centronucleation index, regression of fibrosis and inflammation, and significant improvement of global muscle force compared to untreated dystrophic mice. Transcriptomic analysis of the GA muscle further supported the efficacy of the combined therapy, showing enhanced transcriptomic correction when µ-dys was combined with trehalose compared to µ-dystrophin treatment alone. Notably, the inclusion of trehalose in the combined therapy promoted transcriptomic normalization across various signaling pathways and biological processes, including immune/inflammatory response, calcium homeostasis, cell death pathways, lipid metabolism, endocytosis, and the mTOR pathway. It is however noteworthy that trehalose alone did not have a striking effect on transcriptomic correction, suggesting that it may be acting synergistically with µ-dys in the combined therapy set-up. Additionally, in models of neurodegenerative diseases, trehalose has been shown to modulate the intestine microbiota, oxidative stress, and inflammation (Khalifeh et al., 2021), all which are dysregulated in DMD. Therefore, it is possible that the beneficial effects of trehalose observed in this study are mediated through one or more of these pathways in addition to its action on the endolysosomal system.

One intriguing observation in this study is the partial resolution of lysosomal damage achieved through the expression of the µ-dys protein. This contrasts sharply with the high-level correction of lysosomal damage seen with gene therapy in the LGMDR5 model, where the expression of the complete gamma-sarcoglycan protein is achieved. This discrepancy is particularly notable given the pathophysiological similarities between these closely related muscular dystrophies. A key difference between the two gene therapies is the restoration of the full-length gamma-sarcoglycan protein in the LGMDR5 model, compared to the expression of a shorter, partially functional form of dystrophin (µ-dys), which lack most of the spectrin-repeat rod domain, in the *Dmd^mdx^* model. This suggests that the missing portion of µ-dys may play a crucial role in regulating lysosomal functions in skeletal myofibers. Detection of lysosomal damage in the mouse model for LGMDR5 indicates that this damage might not be specific for DMD. However, both DMD and LGMDR5 are muscular dystrophies that affect the dystrophin-associated glycoprotein complex (DAGC). Whether lysosomal damage occurs in non-DAGC related muscular dystrophies and myopathies remains to be determined.

In summary, our investigation highlights a previously overlooked critical role of lysosomal damage in the pathophysiology of DMD. In the *Dmd^mdx^* mouse model, this damage is only partially corrected by µ-dys gene therapy. We demonstrate that this damage can be alleviated with trehalose, especially when used in combination with gene therapy to restore a functional form of dystrophin expression. The combination of the restoration of dystrophin expression and the correction of lysosomal damage may offer a new therapeutic perspective in DMD.

## Materials and Methods

### Patients’ biopsies

Open muscle biopsies were obtained for diagnostic purposes from the quadriceps (patient 1) or the deltoid muscles (patients 2, 3, 4 and 5) (see biopsies’ details in **Supplementary Table S1**). The muscle location was not available for patient six. All patients consented for usage of the biological samples for research purposes. The specimens were snap frozen in isopentane cooled in liquid nitrogen for histoenzymology and immunohistochemistry. Transversal cryosections were stained using standard histological and histoenzymological methods. Histologically normal muscles served as controls. The genetic confirmation was done by sanger sequencing.

### Animal experimentation and tissue sampling

Wild-Type (WT) (C57BL/10J and C57BL/6J) mice, *Dmd^mdx^*(C57BL/10ScSn-Dmdmdx/J) and *Dmd^mdx-4Cv^* (B6Ros.Cg-Dmdmdx-4Cv/J) mice were supplied by Charles Rivers Laboratories, Miserey-France. Animals were housed in cages with regular 12-hour day/12-hour night cycles, in a specific-pathogen free (SPF) barrier facility and provided with water and food ad libitum. All animal experimentation procedures were conducted according to French and European guidelines for the care and use of animals for experimental purposes. All procedures were approved by ethical committee CEEA-51 and the French Ministry of Higher Education, Research, and Innovation under APAFIS numbers #19736 (DAP2018-024-B) and #35896 (DAP2022-004-C). Only male mice were used for our studies. The high-fat cholesterol-rich diet was supplied by SAFE (260HF diet supplemented with 20g cholesterol/kg). Trehalose was supplied by sigma (ref. T9449) in powder form and dissolved in the mice’s drinking water at a 2% concentration, which was changed twice a week. For rAAV injections, 100 µL of viral vectors diluted in saline solution were injected intravenously into the tails of the mice. For blood sampling, mice were anesthetized with isoflurane, or a mixture of ketamine (100mg/kg) / xylazine (10mg/kg), and blood was collected from the retro-orbital vein on dry capillaries and centrifuged at 10000 rpm for 10 minutes. Serum was then collected and stored at -80°C until analysis. Mice were sacrificed by cervical dislocation at the end of the studies, and tissues were either dissected, mounted transversely on corks and frozen in cooled isopentane, or frozen directly in liquid nitrogen. Samples were stored at -80°C until molecular and histological analyses.

### Dogs’ biopsies

The dogs included in this study were housed and cared for according to current EU regulations. Muscle biopsies were sampled in the context of a project authorized by the the ethical committee ComEth ANSES-EnvA-UPEC (national registration number #16) under the approval number #11-01-11/06. Muscle biopsies from GRMD and WT littermates were surgically sampled from the tibialis cranialis muscle under general anesthesia (IV propofol induction, maintenance isoflurane in 100 % O_2_) with an adapted analgesia (IV morphine). The muscle biopsies were immediately snap-frozen in isopentane cooled in liquid nitrogen, and subsequently stored at -80°C until cryosectioning.

### Functional Evaluation

An escape test was performed to measure the overall muscular force of mice according to standard procedures. Briefly, mice were placed in a 30 cm-long tube with both ends open, and their tails, which were connected to a force transducer, were pinched to induce escape. The force exerted was then measured by the detector. The procedure was performed 5 times, and the average of the 5 measured forces was normalized to the mouse’s weight.

### Transgene constructs and rAAV production

The µ-dys construct has been previously described (Bourg et al., 2022). The µ-dys coding sequence was inserted into a transcription cassette under the control of the muscle promoter Spc5.12, flanked by our optimized sequences of the inverted-terminal repeats (ITR) (referred to as ITR-GNT), and encapsulated in a serotype 9 AAV. The γ-sarcoglycan (*SGCG*) construct has been previously described (Israeli et al., 2019). The sequence of the human copy of *SGCG* was inserted into a transcription cassette under the control of the desmin muscle promoter, flanked by the WT ITR sequences (ITR-2), and encapsulated in a serotype AAV 9 vector.

All rAAV in our studies were produced in HEK 293-T cells using the triple transfection protocol. For that, HEK 293-T cells were grown in suspension (seeded at 0.25×106 cells/mL) under agitation (37°C, 5% CO2) in Freestyle F17 medium (Life technologies, ref. A13835-01), supplemented with GlutaMAX-I at 4nM (Life technologies, ref. 35050-038), Pluronic F-68 at 0.08% (Life technologies, ref. 24040-032) and Anti-clumping agent (Life technologies, ref. 01-0057AE). Three days after seeding, cells were transfected with three plasmids encoding adenovirus helper proteins (pAd Helper), AAV Rep and Cap proteins, and the transgene expression cassette flanked by ITRs. Three days after transfection, cells were harvested, chemically lysed, treated with benzonase (Millipore) and filtered. Viral capsids were purified by affinity chromatography, formulated in sterile PBS and stored at -80°C.

Titers of rAAV vector stocks were determined using real-time PCR (qPCR). Viral DNA was extracted using MagNA Pure 96 DNA and the NA small volume viral kit (Roche Diagnostics) according to the manufacturer’s instructions. PCR was performed with Absolute ROX mix (Taqman, Thermo Fisher Scientific), using ITR-specific primers (List in Supplementary **Table S2**).

### Cell Culture

Immortalized human myoblasts were generated from a healthy donor (C25 cell line) at the Myoline platform (Institue of Myology, Paris). Immortalized myoblasts were grown in skeletal muscle growth medium (Promocell, ref. C-23060), supplemented with a supplement mix (Promocell ref. C-39365), 15% fetal bovine serum (Gibco 10270-106), 1% GlutaMAX (Thermo Scientific, ref. 35050061) and 1% gentamycin (Life technologies, ref. 15750-037). For myotube generation, myoblasts were grown until confluency, and growth medium was then replaced by a differentiation medium (Promocell, ref. C-23061), and cells were differentiated for 6 to 7 days before read-outs. For lysosomal injury assays, myoblasts were plated on a clear-bottomed microscopy dish (Ibidi, ref. 81158) (∼60,000 cells/cm^2^) or in 6-well plates and differentiated into myotubes. Myotubes were then treated with a solution of L-Leucyl-L-leucine methyl ester (LLOMe) (Santacruz, ref. sc-285992) at a final concentration of 2.5 mM in the medium and incubated for the indicated time at 37°C, after which the medium was collected, and the cells were either pelleted and frozen at -80°C for western blotting or fixed for immunofluorescence assays.

For immunofluorescence, cells were briefly washed 2 times with HBSS buffer (Hanks’ Balanced Salt Solution, calcium & magnesium, Gibco), fixed with 4% ice-cold paraformaldehyde (PFA) solution (Thermo Scientific, ref. 28906) for 15 min, then washed with PBS 3 times and permeabilized for 10 min with 0.25% Triton X-100 (Sigma). Immunofluorescence was performed according to protocol described below.

For LGALS3 measurement in the media, media was collected after LLOMe treatment and frozen at -80°C. An ELISA for human LGALS3 (Invitrogen, kit BMS279-4) was performed according to the manufacturer’s instructions.

Dextran pH probing was performed as described before (Fernandez-Mosquera et al., 2019). Briefly, myotubes were loaded with a with mix of Dextran Tetramethylrhodamine (10 000 MW, Invitrogen D1817) and Dextran Oregon Green (10 000 MW, Invitrogen D7171) at a final concentration of 100 µg/mL and incubated for 6h at 37°C. Cells were then washed with HBSS and incubated overnight in fresh media (chasing of the dextran into the lysosomes). Later, LLOMe was diluted in the media at 2.5 mM concentration for 30 min, after which the cells were washed twice with HBSS, fixed in 4% ice-cold PFA and mounted in DAPI-Fluoromount-G (Southern Biotech, ref. 0100-20).

### Molecular Analysis

#### Viral genomes copy number quantification

Genomic DNA (gDNA) was extracted from frozen muscle and organ samples or sections. Briefly, samples (frozen muscle and tissue sections) were lysed using the Beadmill (FisherBrand) machine with RIPA solution (ThermoFisher Scientific, ref. 89901) for 40 seconds (speed =6 m/s), then centrifuged for 10 minutes (2000g, 4°C). DNA was later extracted from the recovered supernatant with the NucleoMag Pathogen extraction kit (Macherey-Nagel, ref. 744210.4) using the KingFisher machine (ThermoFisher Scientific, ref. 5400640). DNA concentrations were quantified by Nanodrop (NanoDrop™ 8000) and stored at - 20°C. For VCN analysis, multiplex channel reactions were performed with the digital droplet PCR (ddPCR) Probe system. Extracted DNA (50 ng) was used with the following PCR amplification conditions: initial denaturation at 95°C for 10 minutes, followed by 40 cycles of denaturation at 94°C for 10 seconds, annealing at 60°C for 10 seconds, and transcript extension at 72°C for 1 minute and 30 seconds. Following this, fluorescent dye stabilization was performed at 98°C for 10 minutes. VCN values were calculated using the ratio with the *Rplp0* gene amplification. For this analysis, multiplex channels were analyzed, depending on the type/concentration of probe. Primers and probes that were used are listed in **Supplementary Table S2**. The analysis was performed using the Quantasoft analysis pro software.

#### RNA isolation and RT-qPCR

Total RNA was extracted from frozen muscle samples or sections. 400μL of Nucleozol solution (Macherey-Nagel ref.740404) were added per sample, and the tissues were lysed using the BeadMill 24 (Fisherbrand) (6m/s – 40s). Next, 160μL of DNAse/RNAse free water were added, and the tubes were mixed vigorously and incubated at RT for 10 minutes. Samples were then centrifuged (1000g, 15min, 20°C) and total RNA was extracted from the supernatant using magnetic beads (Macherey-Nagel, Ref. 744503.12) with the Ideal-32 system (ID solutions). RNA samples were then treated with DNAse (30min ,37°C) (TURBO DNA-free kit, ThemoFisher Scientific), and 1 µg of RNA per sample was used for reverse transcription reaction using the RevertAid H Minus First Strand cDNA synthesis kit (ThermoFisher Scientific, K1632) according to the manufacturer’s instructions. The cDNA samples were then diluted (1:9) and analyzed by qPCR using a LightCycler 480° (Roche), with either SYBR Green PCR assays (ThermoFisher Scientific, ref.4309155) or taqman gene expression assays (ThermoFisher). The primer sequences used are shown in **Supplementary Table S2**. Each reaction was performed in duplicate, and the Ct values for *Rplp0* were used to normalize gene expression using Livak’s secondary derivation method (Livak & Schmittgen, 2001).

#### Protein isolation and western blotting

Frozen tissues were cut using a cryostat into 10 µm sections, and lysed with RIPA lysis buffer (Thermofisher Scientific, Ref-89901) using the the Beadmill 24 (Fisherbrand) for 40 seconds at 6m/s. Samples were then centrifuged for 10 minutes (2000g, 4°C), after which the supernatant was collected and treated with DNAse (1:1000, Millipore) and a protease inhibitor cocktail (Roche) for 45 minutes. Total protein concentration was measured using the Pierce® BCA Protein Assay kit (Thermo Fisher Scientific, ref. 23225). Equal amounts of protein (between 20 and 30 μg) were loaded and separated on NuPAGE pre-cast gels (Thermo Fisher Scientific). Proteins were transferred to nitrocellulose membranes using the iBlot 2 Dry Blotting system (Thermo Fisher Scientific), and total protein quantification was performed using the Revert Total protein Stain kit (Li-Cor). Membranes were later incubated with a blocking solution (Odyssey Blocking Buffer, Li-Cor) for 1h at room temperature (RT). Primary antibodies incubation was done overnight at 4°C (dilutions and details of primary antibodies are presented in **Supplementary Table S3**). Membranes are then washed in a TTBS solution 3 times and incubated with secondary antibodies (1:10 000, Listed in **Supplementary Table S5**). Membranes are washed again with TTBS and scanned using the Odyssey CLx scanner (Li-Cor). Images were analyzed using Image Studio Lite (ver. 5.2), pixels of selected bands were quantified, and the signal was normalized to either reference protein (GAPDH) or to total protein staining.

#### Lipidomic analysis

Muscle sections were weighted and lysed in PBS, then diluted to 5 mg/mL. Shotgun lipidomic analysis was performed by Lipotype (Dresden, Germany). This technique involves automated sample extraction, automated direct sample perfusion and high-resolution mass spectrometry to capture lipid class-specific internal standards. Briefly, lipid extraction was performed using chloroform and methanol (Surma et al., 2021). Samples were spiked with lipid class-specific internal standards prior to extraction. After drying and resuspension in the mass spectrometry acquisition mixture, the lipid extracts were subjected to mass spectrometry analysis. Mass spectra were acquired on a hybrid quadrupole/Orbitrap mass spectrometer equipped with an automated nano-flow electrospray ion source in positive and negative ion modes. LipotypeXplorer’ software is used for lipid identification on mass spectra. Data processing and analysis are carried out using the Lipotype information and management system and the LipotypeZoom web software. **Quantification of free muscle cholesterol:** Muscle sections were weighed (between 10-20 mg) then lysed using a mix of chloroform/Isopropanol/NP-40 (7 : 11 : 0.1) for 1 minute with the Beadmill 24 (Fisherbrand). Samples were then centrifuged at 15,000 g for 10 minutes, and supernatants were transferred to new tubes, dried at 50°C and centrifuged under vacuum for 30 min. The pellet is then re-suspended in cholesterol quantification kit buffer (Invitrogen, ref. A12216) at a concentration of 20 μg/μL, and vortexed vigorously. Free cholesterol quantification was performed in duplicate using the Amplex Red Cholesterol kit (Invitrogen, ref. A12216), following the manufacturer’s instructions. Cholesterol concentrations were normalized to tissue weight for each sample.

#### RNA sequencing

Essentially as described previously (Vu Hong, 2023), RNA was extracted from muscle sections, and purified RNA samples were first checked for RNA quality using the Bioanalyzer 2100 (Agilent). Samples with an RNA integrity index greater than 8 were then used for RNA sequencing (Genewiz). The Stranded Total RNA Library Prep Kit (Illumina) was used to create sequencing libraries, which were sequenced according to Illumina protocol on the NovaSeq instrument (Illumina), yielding around 20 million paired end reads per library. Paired reads were filtered and subjected to quality-control using fastp. They were then mapped onto the GRCm38/mm10 genome using HISAT2 and count tables generated using htseq-count Differentially expressed genes (DEGs) were identified using the DESeq2 R package. Pathway analysis was carried out in R-Studio (version 4.0.4) using either over-representation methods with Gene Ontology or functional class scoring with Gene Set Enrichment Analysis (GSEA). GSEA was performed using the *clusterprofile* R package.

#### Serum biomarkers analyses

Quantification of creatine kinase (CK) in serum was performed using a FUJI DRI-Chem nx500 analyzer (DMV Imaging). 10 μL of diluted serum was deposited on FUJI DRI-CHEM CPK-PIII plates (DMV Imaging, ref.15809463), and CK concentrations were automatically generated in international units per liter of serum (U/L). Fragments of the myofibrillar structural protein myomesin-3 (MYOM3) have been identified as being abnormally present in the sera of dystrophic animal models (Rouillon et al., 2015). To quantify these fragments in mouse serum, a Meso Scale Discovery (MSD) enzyme-linked immunosorbent assay (ELISA) was performed. Anti-MYOM3 polyclonal capture antibody (Proteintech Lab, ref. 17692-1-AP) was coated to the wells of a multi-array plate containing electrodes (MSD, ref. L15XA-3) (overnight, 4°C). The plate was then washed 3 times with a washing solution (PBS + 0.05% Tween 20), and the wells were saturated with a 3% bovine serum albumin (BSA) solution (1h, RT). After 3 washes, diluted sera were added (1:50 dilution in 1% BSA solution) and the plate was incubated for 2 h at room temperature under rigorous agitation. A standard range of MYOM3 peptide was also used to calculate concentrations. Revelation was carried out using a mouse monoclonal antibody (Anti-51r20, developed by Proteogenix©) coupled to a sulfo-tag (Mesoscale, ref. R31AA-2). The plate was incubated for a further 2 hours under agitation, then washed 3 times. Finally, 150 μL of MSD-Gold reading buffer (MSD, ref. R92TG-2) was added and the electro-chemiluminescence was read using the MESO QuickPlex SQ 120 (MSD). MYOM3 concentrations were calculated using the MYOM3 standard range and expressed in pg/μL.

### Single Myofiber isolation and immunostaining

The two Flexor Digitorum Brevis (FDB) muscles were surgically isolated from euthanized *Dmd^mdx-4Cv^* and C57BL/6J male mice (6 weeks old), rinsed in PBS and placed in a digestion solution (collagenase 2.5U/mL in DMEM, Sigma-Aldrich, United States, 11088793001) for 1h30 at 37°C on a rotated wheel. The collagenase solution was removed by fiber sedimentation (10 min, RT) and the isolated muscle fibers were transferred in 2 mL tube and fixed with an ice-cold 4% PFA solution for 20 minutes. Fixed myofibers were then washed 3 times with PBS (10 min – RT) and permeabilized with 0.5% Triton-X-100 (Sigma) for 10 minutes. Afterwards, 10% goat serum (GS) (Agilent, ref. X090710) was used for blocking (30 min – RT), before a second blocking with ‘mouse-on-mouse’ blocking solution (Invitrogen, ref. R37621). Primary antibodies (listed in **Supplementary table S4**) were incubated overnight at 4°C on an inclined agitator. Myofibers were then washed 3 times with a Tween-Tris-buffered saline (TTBS) solution (10 min – RT), before the incubation with secondary antibodies (Listed in **Supplementary Table S5**, 1:200 dilution). Finally, myofibers were washed 3 times with PBS and mounted on glass slides with a DAPI-Fluoromount-G mounting medium (Southern Biotech, ref. 0100-20).

Stack images were acquired with a LEICA TCS SP8 spectral confocal microscope (Leica, Germany) running LASX software (Leica, Germany), using a HC PL APO CS2 20X 0.75 NA dry objective. A stack image was taken every z=0.3 µm, and 3D myofibers images were reconstructed using Imaris software (version 9.9.1). All myofibers images were treated similarly, and LAMP2 sport analyses was performed using spot modeling function (average spot diameter = 1µm).

### Histological studies and image analyses

For all histological studies, muscles were dissected, fixed on corks and frozen in isopentane cooled in liquid nitrogen, then stored at -80°C. Transverse cryosections (8μm thick) were prepared from frozen muscles, air-dried, and stored at -80°C.

#### Histochemistry

**H&E staining** was carried out in accordance with standard procedures. This staining combines nuclear staining with hematoxylin and cytoplasmic staining with eosin. Analysis of myofiber size distribution and centronucleation index was performed as previously described in (Reinbigler et al., 2022).

**Oil Red O** staining identifies hydrophobic lipids in the liquid phase. The colored body, called Lysochrome, moves from the solvent in which it is found to the tissue lipid, and the lipid droplets are colored red and the nuclei blue.

**Sirius Red staining** was used to assess tissue fibrosis, combining red staining for collagen fibers with yellow staining for cytoplasm. Briefly, slides were treated with acetone for 1 hour, fixed in formalin for 5 minutes, then incubated in Bouin’s solution for 10 minutes. Slides were then rinsed twice with water and stained in Sirius Red solution (Sigma, ref. 365548) (diluted in 0.1% picric acid) for 1 hour, washed with water and dehydrated with ethanol (70% 1 min, 95% 1 min, 100% for 2 min) and then cleared in a xylene bath (2 washes for 1 min). Slides were mounted in mounting medium and air-dried.

All bright-field histological labeling images were digitized using an Axioscan Z1 slide scanner (Zeiss, Germany) under a Zeiss Plan-Apochromat 10X/0.45 M27 dry lens (ZEISS, Germany). Tiled scanner images were reconstructed using ZEN software (ZEISS, Germany) (pixel size = 0.45 µm). QuPath (version 0.4.3) software was used for muscle fibrosis quantification from sirus red staining. For each muscle scan, a small artificial neural network was trained to classify positive and negative fibrotic pixels, and subsequently used to quantify fibrotic areas.

### Immunofluorescence

#### Galectin-3 and all co-immunostainings of LAMP2

Slides were fixed in ice-cold 4% PFA (Thermo Scientific, ref-28906) for 3 minutes and washed 2 times in PBS, then permeabilized in an ice-cold methanol gradient (10 min 70%; 10 min 95%, 20 min 100%). After 3 washes with PBS, a 10% goat serum (GS) solution (Agilent, ref. X090710) was used for blocking (RT, 30 min). A second ‘Mouse-on-Mouse’ blocking solution (Invitrogen, ref. R37621) was also used to block endogenous binding to mouse immunoglobulins (RT, 30min), for all immunostainings with mouse primary antibodies on mouse muscles. Slides were then incubated with primary antibodies at 4°C overnight (diluted in 1% GS, see **Table S3** for specific dilutions). Next, slides were washed in TTBS solution 3 times, before incubation with secondary antibodies for 2h at 4°C (Molecular Probes Alexa Fluor secondary antibodies were used, 1:200 dilution, see **Supplementary Table S5**). Finally, the slides were washed in PBS solution 3 times for 5 minutes, rinsed in milliQ water and mounted with DAPI-Fluoromount-G mounting medium (Southern Biotech, ref. 0100-20).

#### Other immunostainings

Slides were fixed for 10 minutes in 4% methanol-free ice-cold PFA solution (Thermo Scientific, ref. 28906), permeabilized in 0.1% Triton for 5 minutes and blocked for 45 minutes in 10% goat serum solution (Agilent, ref. X090710). For staining using primary mouse antibodies, an additional 30-minute blocking was done with ‘mouse on mouse’ solution (Invitrogen R37621). Next, slides were incubated with the primary antibody solution at 4°C overnight (diluted in 1% GS, references and dilutions are listed in **Supplementary Table S4**). Slides were then washed 3 times in PBS and incubated 2h at 4°C with a secondary antibody solution (Molecular Probes Alexa Fluor secondary antibodies were used, 1:200 dilution, listed in **Supplementary Table S5**). Finally, the slides were washed in PBS solution 3 times for 5 minutes, rinsed in milliQ water and mounted with DAPI-Fluoromount-G mounting medium (Southern Biotech, ref. 0100-20) or Fluoromount-G medium (Southern Biotech, ref. 0100-01).

#### Detection of free cholesterol by filipin staining

Free cholesterol was detected using the fluorophore filipin (Sigma, Ref. SAE008). Filipin was incubated with the secondary antibody solution at a concentration of 100μg/mL, along with Syto13 (Invitrogen, S7575) for nuclear staining.

### Image acquisition

Once the slides have been dried, they were imaged with a LEICA TCS SP8 spectral confocal microscope (Leica, Germany) running LASX software (Leica, Germany), using a HC PL APO CS2 20X 0.75 NA dry objective. Acquired images were processed with Fiji ImageJ for final figures.

For dystrophin, IgG and CD11B staining, images were digitized using the Axioscan Z1 slide scanner (Zeiss, Germany) under a Zeiss Plan-Apochromat 10X/0.45 M27 dry lens (Zeiss, Germany) and using an ORCA-Flash4.0 CMOS digital camera (Hamamatsu, Japan). Tile scanner images were reconstructed with ZEN software (ZEISS, Germany) (pixel size = 0.65 μm).

Image Analyses:

#### Myofiber segmentation

For images with laminin or wheat germ agglutinin (WGA) labeling for the membranes, the Cellpose2 *cyto2* model (Stringer et al., 2021) was fine-tuned on manually myofiber-labeled images (hyperparameters: n_epochs=200, learning_rate=0.05, weight_decay=0.0001). The labeled dataset used in fine-tuning was prepared in such a way that the model can simultaneously segment myofibers and ignore low-quality staining areas. Fine-tuned models were then used to extract myofiber masks. Reconstruction of myofiber masks of whole-scan images was done using the *cellpose* package (Stringer et al., 2021). Reconstructed masks were then converted into Regions of Interest (ROIs) for subsequent quantification (each ROI corresponds to an individual myofiber) using the *Labels_To_Rois.py* FIJI plugin (Waisman et al., 2021). For all images acquired with the confocal, the cellpose2 *cyto2* model was fine-tuned on manually labeled images based on the background noise of the A488 channel.

#### Puncta Quantification

all images from the same experiment or panel were processed uniformly using Fiji ImageJ. Median filters were applied to reduce noise and thresholding was performed to select the puncta to be analyzed. For overlay quantification, the image calculator function ‘AND’ was used. Various parameters of puncta were quantified within each individual region of interest (ROI), where 1 ROI corresponds to 1 myofiber.

#### Overlay analyzes

ImageJ’s JaCOP plug-in was used to analyze colocalization by calculating Manders coefficients. These coefficients indicate the fraction of the fluorescence signal from one channel localized in the fluorescence pixels of a second channel.

### Statistical Analysis

Statistical analysis was performed using Graphpad Prism 10 (GraphPad Software, Inc., La Jolla, CA, USA), R 4.2.2. or Python 3.10. Variances were assumed to be equal; a Shapiro-Wilk test was used to check normality. A Student’s t test was used to compare the mean of two groups. A one-way ANOVA test was used to compare more than two groups when values followed a normal distribution, accompanied by Tukey’s multiple comparison test, otherwise a non-parametric Kruskal-Wallis test with Dunn’s multiple comparisons was used. Error bars on graphs represent standard deviation (SD) unless specified otherwise. Results were considered significant when p-values or adjusted p-values were less than 0.05.

## Acknowledgments

This work was supported by the “Association Française contre les Myopathies” (AFM), and “Institut National de la Santé Et de la Recherche Médicale” (INSERM, FranceRelance). The authors are Genopole’s members, first French biocluster dedicated to genetic, biotechnologies and biotherapies. We are grateful to the “Imaging and Cytometry Core Facility” and to the *in vivo* evaluation, services of Genethon for technical support, to Ile-de-France Region, to Conseil Départemental de l’Essonne (ASTRE), INSERM and GIP Genopole, Evry for the purchase of the equipment.

## Funding

For I.B and S.B, the work was funded by the Association Française contre les myopathies (Translamuscle II (22946)).

## Author contributions

The project was conceptualized by A.J and D.I. A.J conducted most of the experiments and performed data analysis. R.B, L.P, E.L, N.B, J.P conducted experiments. L.V.B, A.M and N.D conducted and supervised animal experiments. I.B, S.B, M.T.B and T.E provided muscle biopsies and performed histological experiments. A.V.H and D.S performed computational analyses. The original draft was written by A.J. I.R and D.I reviewed and edited the manuscript; all other authors reviewed the manuscript.

## Competing interests

A.J and D.I are co-inventors on filled European patents (EP23194414.1 and EP23194612.0) related to this study.

I.R is co-founder and chief scientific officer of Atamyo Therapeutics. All other authors declare they have no competing interests.

## Data and materials availability

All data are available in the main text or the supplementary materials. RNA sequencing data will be made available publicly upon publishing.

## Supplementary Materials

**Figure S1.**
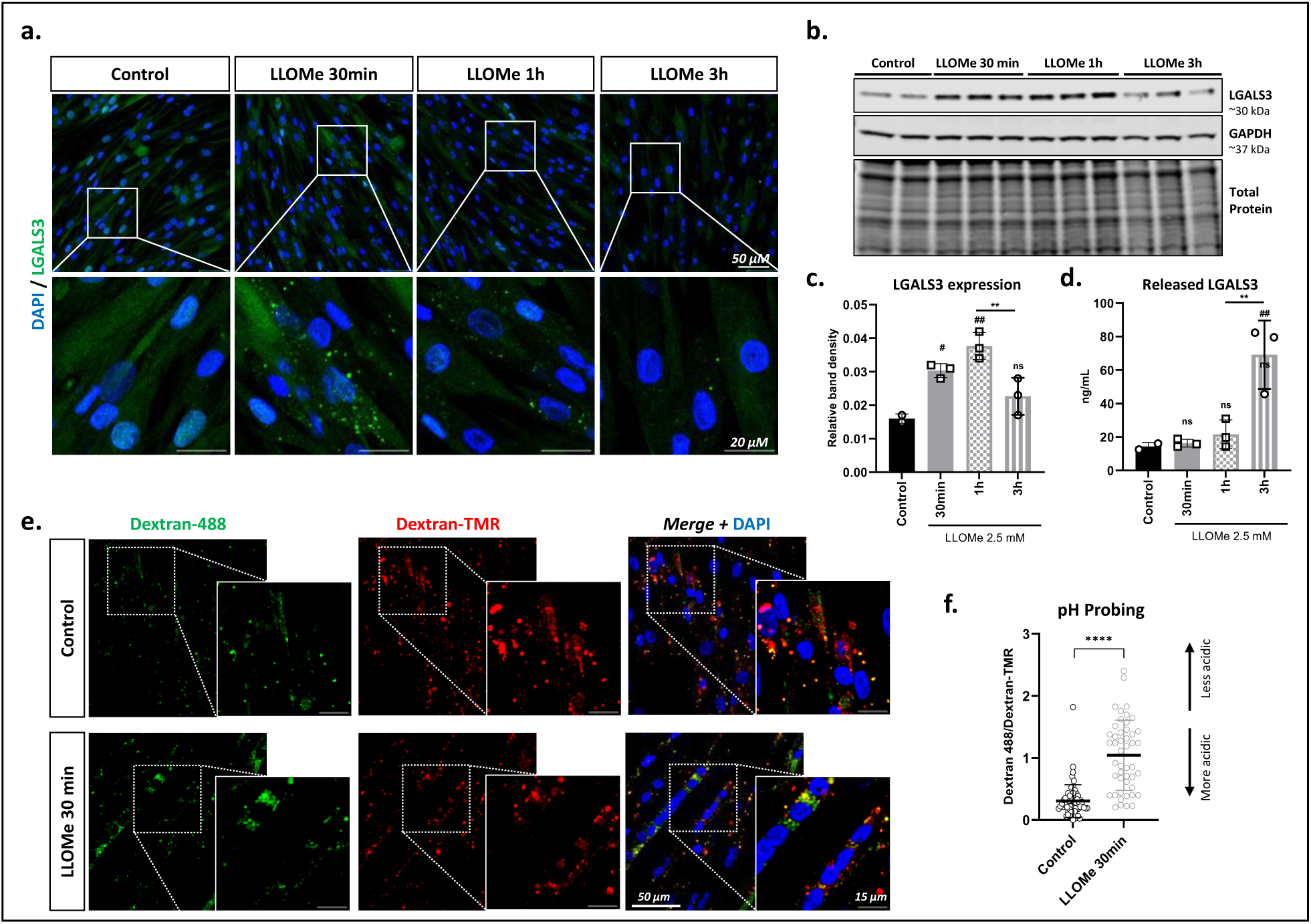
Lysosome membrane permeabilization impairs lysosome function in muscle cells and is detected with LGALS3 puncta assay. (**a**) Representative plane confocal images of differentiated human immortalized myoblasts immunostained for Galectin-3 (green). Cells were treated with LLOMe for 30 minutes, 1h or 3h. (**b**) Western blot analysis of human myotubes for LGALS3, using GAPDH as loading control. (**c**) Column plot shows average ± SD of LGALS3 band intensity normalized to GAPDH. (**d**) ELISA analysis of released LGALS3 in the media of cultured myotubes. (**e**) Representative plane confocal images of human myotubes stained with dextran-Oregon Green and dextran-TMR. Dextran Green is quenched under acidic pH, so increased green:red ratio denotes impaired lysosomal acidification. (**f**) Scatter plot (average ± SD) of Dextran 488/Dextran-TMR ratio for at least 45 myotubes. An ANOVA test with Tukey multiple comparison was used for statistical comparisons of multiple groups. ANOVA p-value #p<0 .05, ##p<0.001, ns=non-significant (compared to Control). An unpaired two tailed t.test was used for 2 groups comparison. T.test p-value ****p<0.0001.

**Fig. S2.**
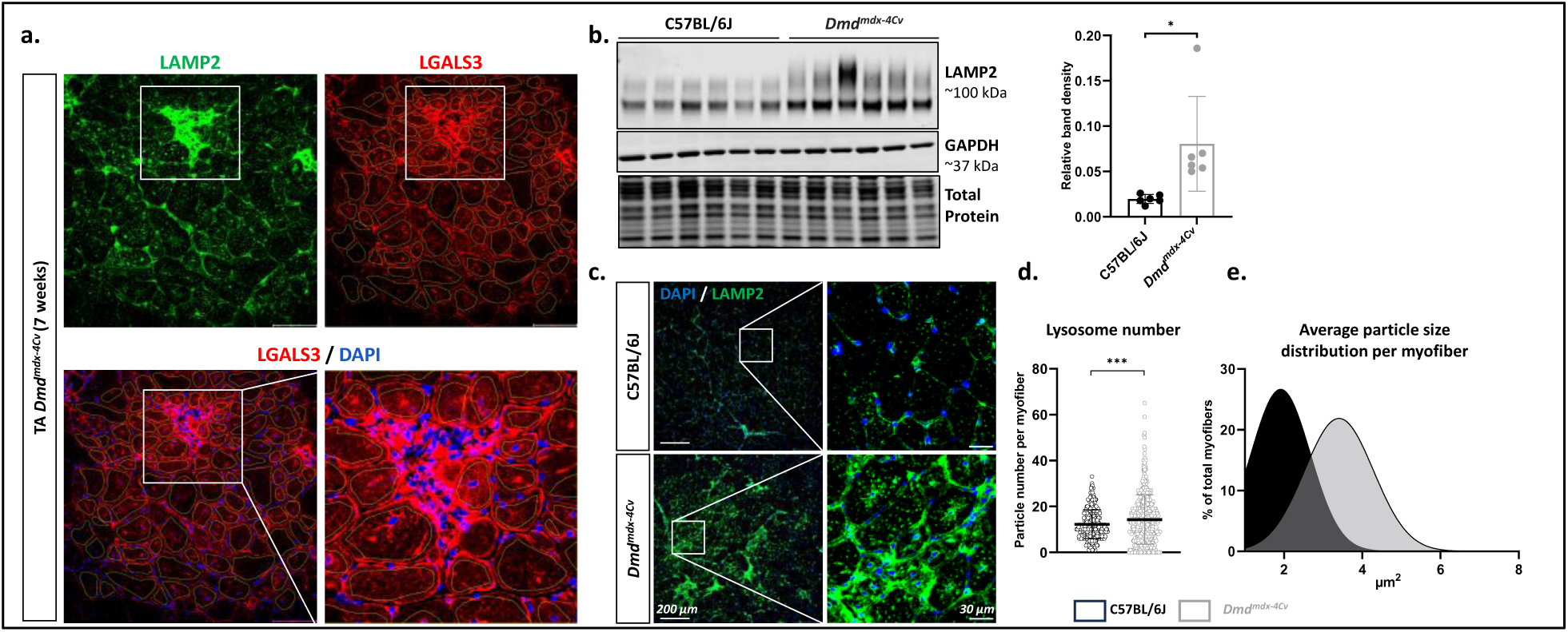
Lysosomal stress in *Dmd^mdx-4Cv^* muscle. (**a)** Confocal images of TA cross sections immunostained for LAMP2 and LGALS3. White boxes represent inflammatory regions in the muscle. Segmentation of regions of interests (used for LAMP2+LGALS3+ quantification) are shown on LGALS3 images, inflammatory regions (with aggregate of nuclei outside the myofibers) are excluded. (**b)** Western blot analysis of TA muscle for LAMP2 and band density quantification of LAMP2, normalized to GAPDH (average ± SD) (n=6). (**c**) Representative confocal images of transversal sections of TA muscle immunostained for LAMP2. (**d**) Scatter plot showing (average ± SD) number of lysosomes per myofiber, detected by LAMP2 immunostaining on transversal sections (>379 myofibers analyzed from n=6). (**e**) Density plot with non-linear fit gaussian curve applied, showing distribution of average spot size per myofiber (>379 myofibers analyzed from n=6).

**Fig. S3.**
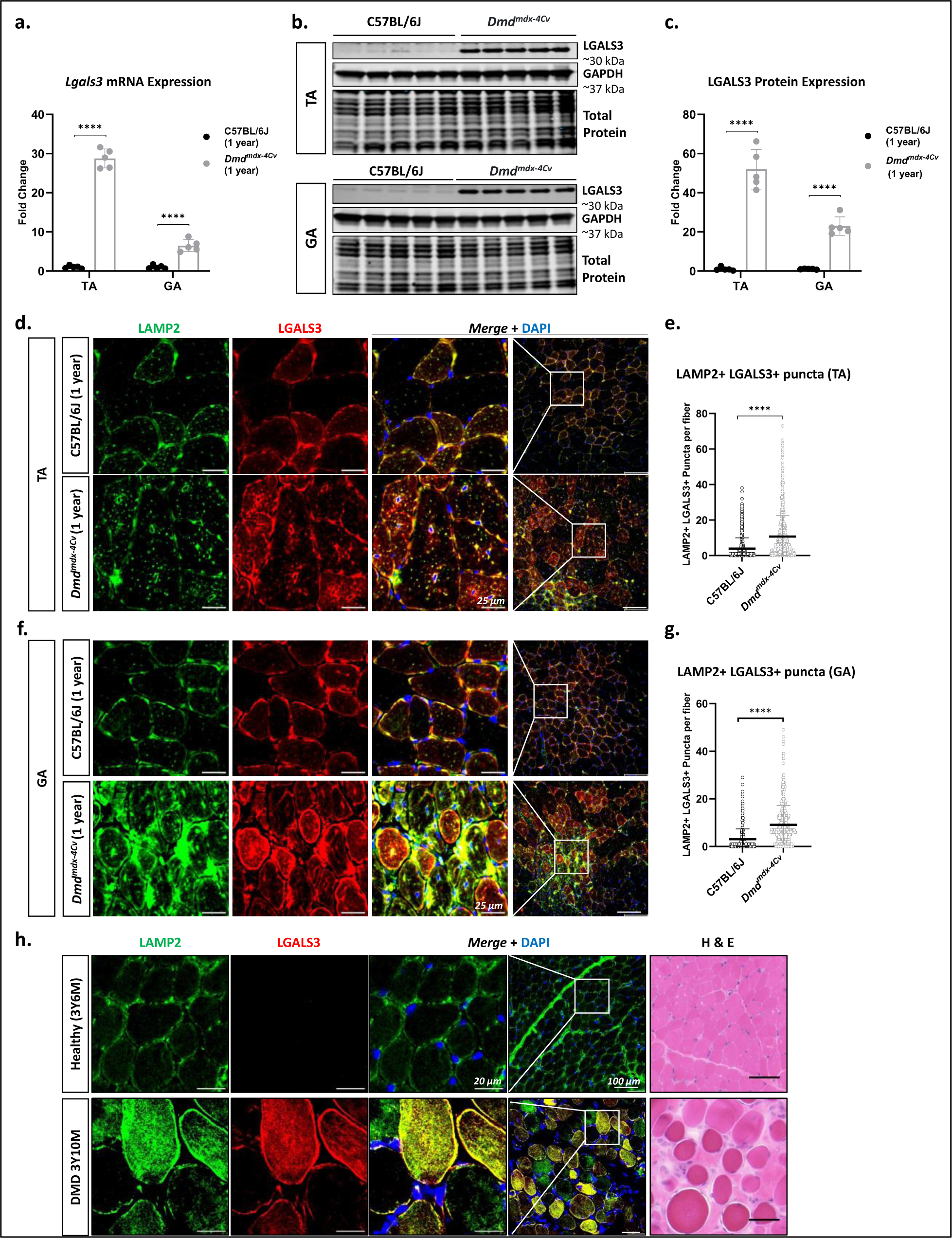
LGALS3 overexpression indicates LMP in old *Dmd^mdx-4Cv^* mice and a DMD patient. **(a)** Relative Lgals3 mRNA expression in the TA, GA, normalized to Rplp0. Data are presented as scatter plots with average ± SD of Fold change relative to WT mice. (**b**) Western blot analysis of TA and GA muscles of 1-year old WT and Dmd^mdx-4Cv^ mice (n=5). (**c**) Band intensity quantification of LGALS3 relative to total protein staining in TA and GA muscles. Scatter plots show average ± SD of Fold change relative to WT mice. (**d-f**) Representative confocal images of transversal sections of TA and GA muscles of 1-year old WT and Dmd^mdx-4Cv^ mice immunostained for LAMP2 (green) and LGALS3 (red). (**e-g**) Quantification of double positive puncta (LAMP2+LGALS3+) in the myofibers of TA (**e**) and GA (**f**) muscles. Data are represented as scatter plots with average ± SD (>838 and >464 myofibers analyzed from n=5). (**h**) Representative confocal images of transversal sections from muscles of a DMD patient and a healthy control immunostained for LAMP2 and LGALS3. Left panels show H&E labeling from the same muscle biopsies. An unpaired two-tailed t.test was used for statistical comparisons of WT and *Dmd^mdx-4Cv^* groups. T.test p-value ****p<0.0001.

**Fig. S4.**
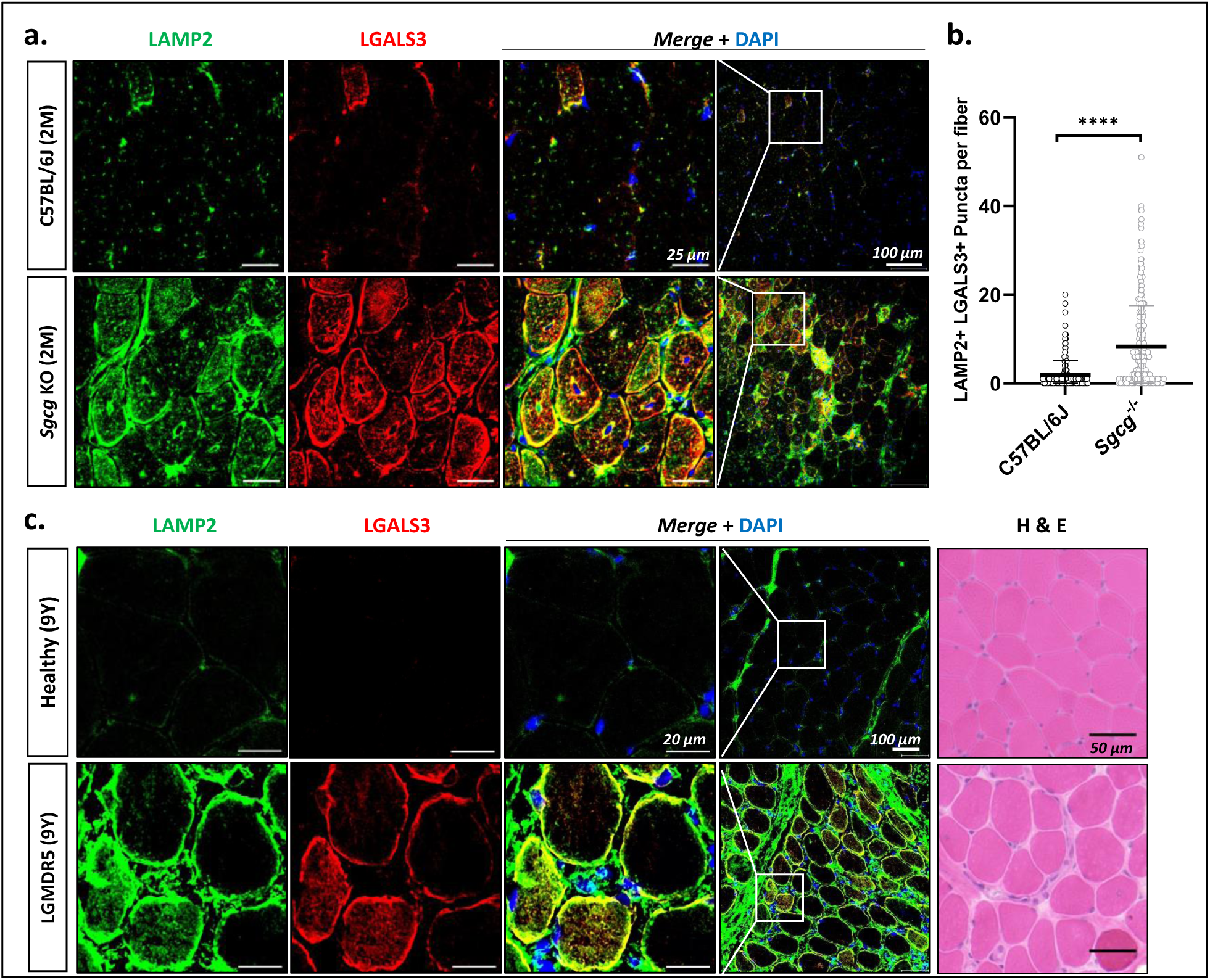
Detection of LMP in LGMDR5 (ɣ-Sarcoglycanopathy). (**a**) Representative confocal images of transversal sections of GA muscles from 2-months old WT and *Sgcg* KO mice, immunostained for LAMP2 (green) and LGALS3 (red). (**b**) Quantification of LAMP2+ LGALS3+ puncta per myofiber in the GA muscle. Data are presented as scatter plot with average ± SD (>338 myofibers from n=5). (**c**) Representative confocal images of transversal sections from muscles of a LGMDR5 patient and a healthy control immunostained for LAMP2 and LGALS3. Left panels show H&E labeling from the same muscle biopsies.

**Fig. S5.**
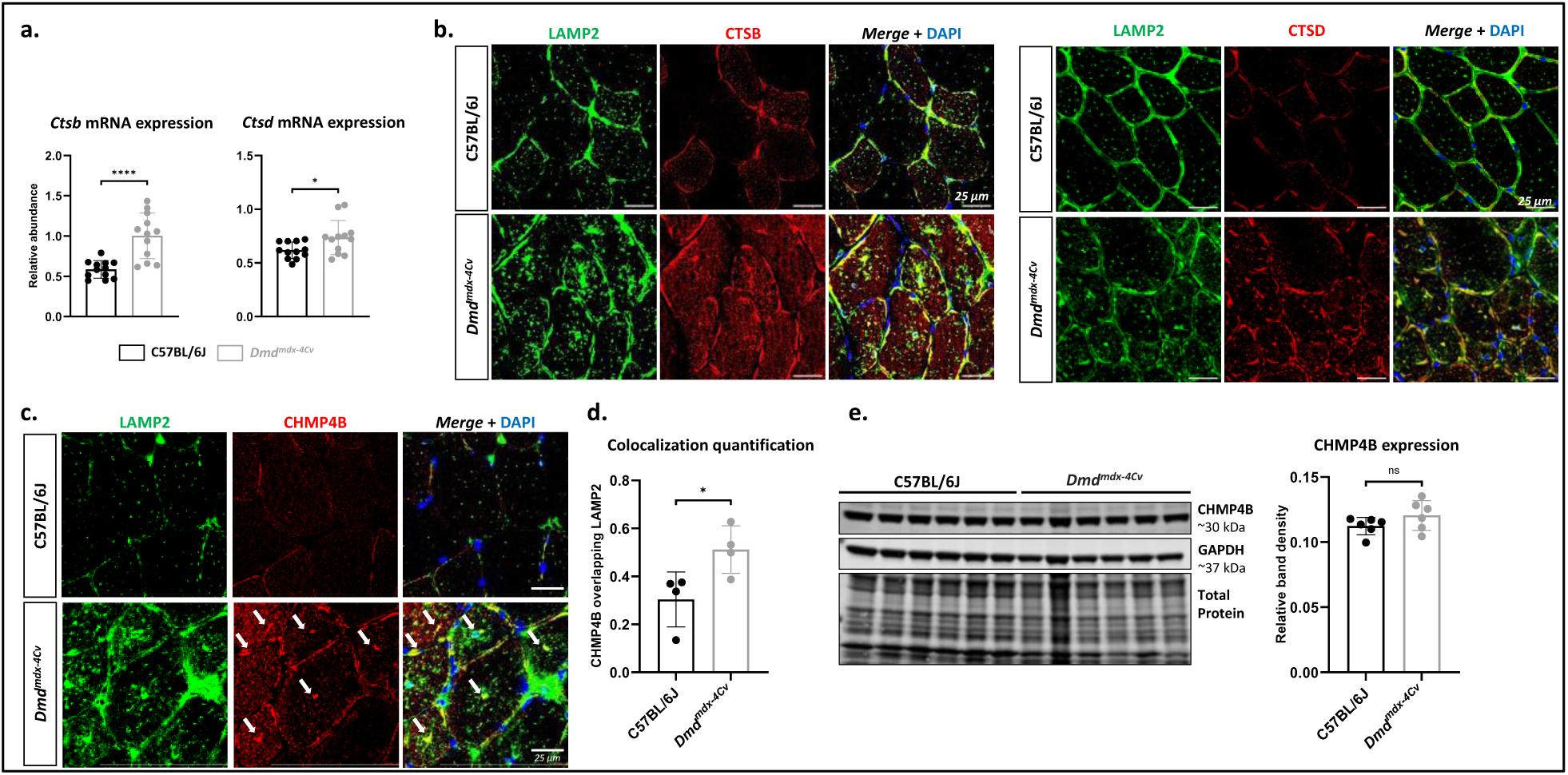
Activation of lysosomal repair pathway in *Dmd^mdx-4Cv^* muscle. (**a)** Relative mRNA expression of Ctsb and Ctsd in the TA, normalized to Rplp0. Scatter plots represent average ± SD (n=12). (**b**) Representative confocal images of TA transversal sections immunostained for Cathepsin B (CTSB) or Cathepsin D (CTSD) and LAMP2. (**c**) Representative confocal images of TA transversal sections immunostained for CHMP4B and LAMP2. (**d**) Quantification of CHMP4B signal overlapping LAMP2 from images of (**c**) (n=4). (**e**) Western blot analysis of TA muscle for CHMP4B, and band density quantification of CHMP4B, normalized to GAPDH (average ± SD) (n=6). An unpaired two-tailed t.test was used for statistical comparisons of WT and *Dmd^mdx-4Cv^* groups. T.test p-value *p < 0.05, ***p<0.001 ****p<0.0001.

**Fig. S6.**
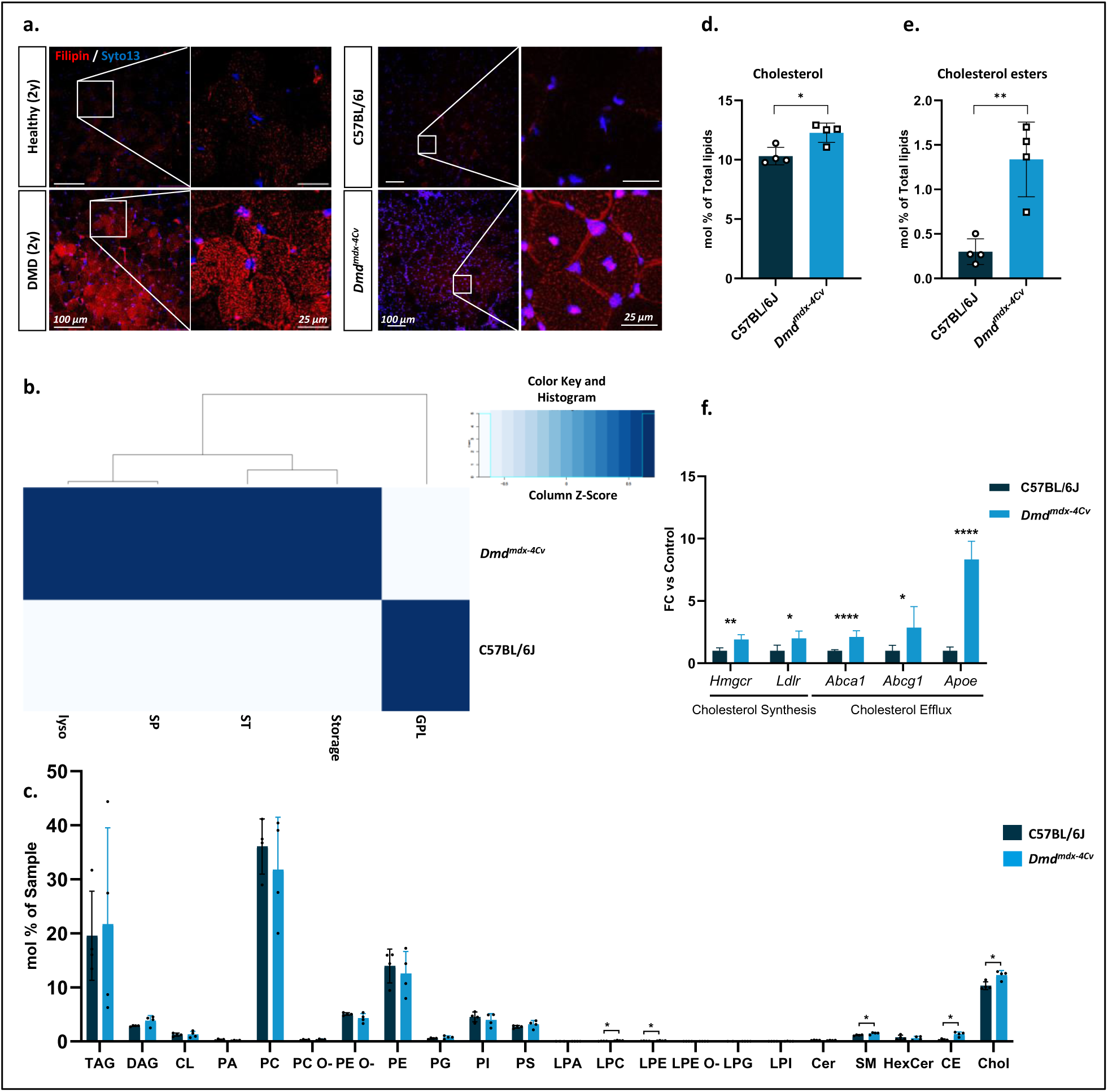
Lipid Perturbations in DMD. (**a)** Representative confocal images showing filipin labeling of transversal muscle cross sections from a DMD patient and healthy control, and a Dmd^mdx-4Cv^ mice and a WT control (GA muscle). (**b**) Heat-Map displaying functional clustering of lipids in GA muscle from 7-week-old Dmd^mdx-4Cv^ mice versus control WT (GPL= glycerophospholipids, ST= sterols, SP= sphingolipids, lyso= lysophospholipids) (n=4). (**c**) Scatter plots showing different classes of lipids (average ± SD). T.Test p-value *p < 0.05 **<0.01. Chol = Cholesterol; CE = Cholesterol Esters; HexCer = hexosylceramide; SM = Sphingomyelin; Cer = Ceramide; LPI = lyso-Phosphatidylinositol; LPG = lyso-Phosphatidylglycerol; LPE O-= lyso-Phosphotidylethanolamine (-ether); LPE = lyso-Phosphotidylethanolamine; LPC = lyso-Phosphotidylcholine; LPA = lyso-phosphotidate; PS = Phosphatidylserine; PI = Phosphatidylinositol; PG = Phosphatidylglycerol; PE O-= Phosphatidylethanolamine (-ether); PE = Phosphatidylethanolamine; PC O-= Phosphatidylcholine (-ether); PC = Phosphatidylcholine; PA = Phosphatidate; CL = Cardiolopin; DAG = Diacyglycerol; TAG = Triacylglycerol. (**d-e)** Scatter plots (average ± SD) showing increase of cholesterol and cholesterol esters in GA muscle of *Dmd^mdx-4Cv^* mice compared to WT controls. T.Test p-value *p < 0.05 **<0.01. (**f**) Relative mRNA expression of different genes involved in cholesterol homeostasis, normalized to Rplp0. Scatter plots represent average ± SD of Fold change compared to WT mice. T-test p-value *p < 0.05 **<0.01 ****<0.0001 (n=5-6).

**Fig. S7.**
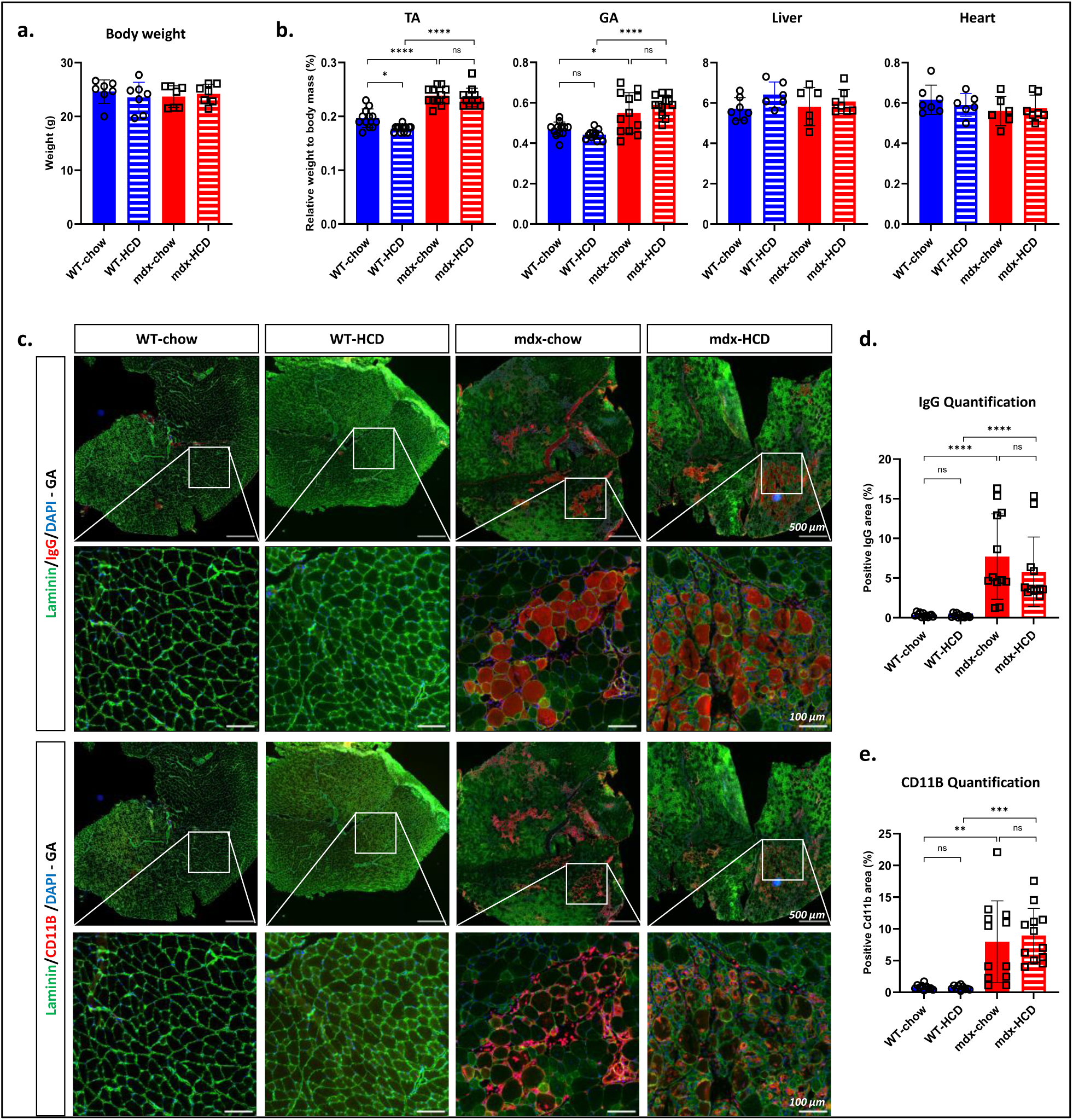
High Cholesterol diet does not affect body and muscle weights in mice and does not increase inflammation or sarcolemma fragility in dystrophic mice. (**a**) Body weigh measurement of mice after 4 weeks of standard or high cholesterol diet (n=7). Data are presented as scatter plot with average ± SD. ANOVA test yields no significant differences across all groups. (**b**) Muscle and tissue weight upon sacrifice, normalized to mice body weight. Data are presented as scatter plot with average ± SD (n=12 for GA and TA, left and right muscles were weighted, n=6 for other tissues). ANOVA p-value *p < 0.05 ****<0.0001. (**c**) Representative images of GA muscle serial cross-sections labeled with Laminin, mouse IgG and CD11b. (**d-e**) Quantification of IgG and CD11b positive areas on whole muscle cross-sections (n=12, 2 cross-sections per mouse were quantified). ANOVA p-value **p < 0.01 ***<0.001 ****<0.0001 ns=non-significant.

**Fig. S8.**
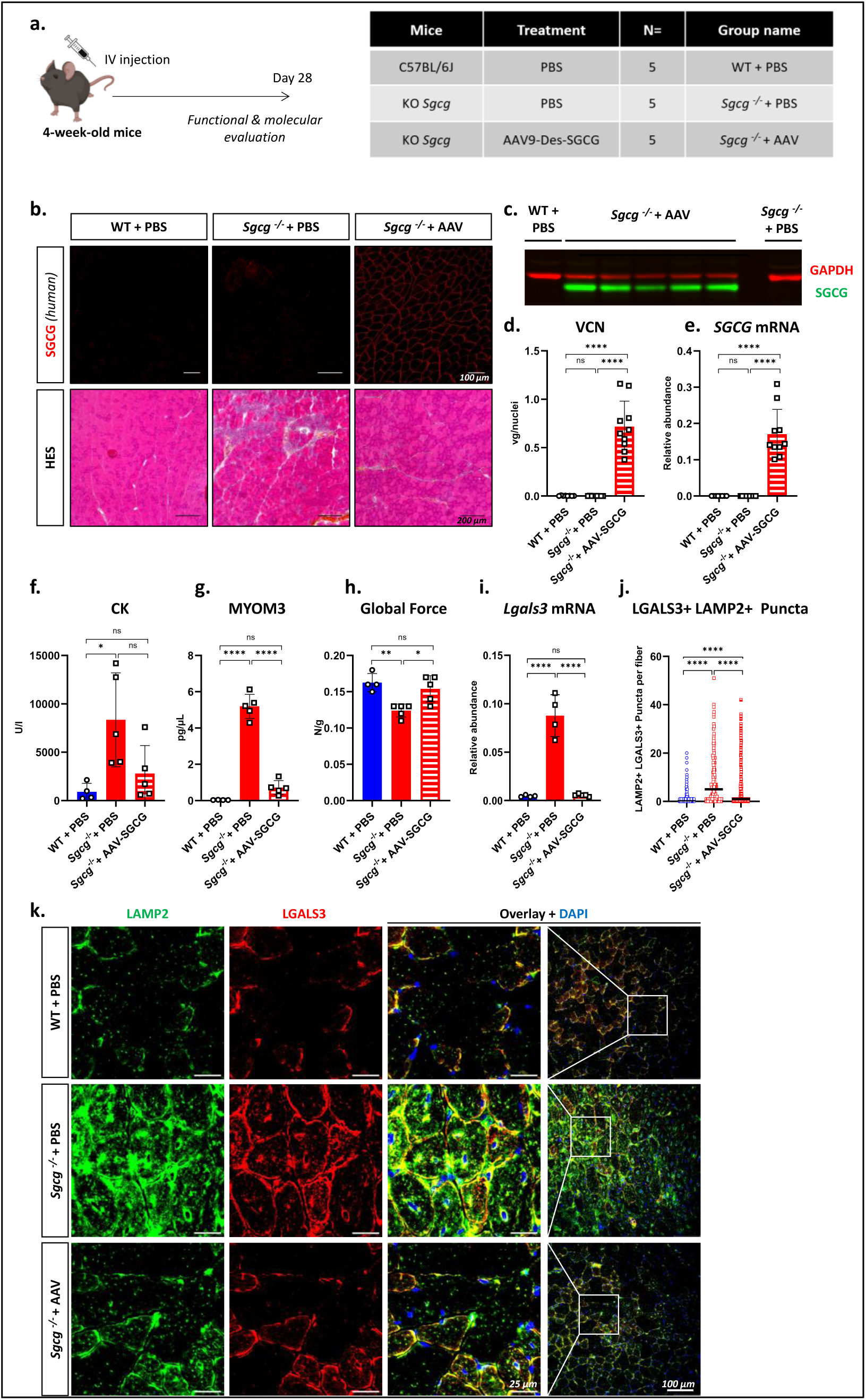
Lysosomal damage is partially corrected by gene therapy in *Sgcg^-/-^* mouse model. **(a)** Schematic representation of study set-up. 4-week-old *Sgcg^-/-^* mice were injected intravenously with a rAAV9 encapsulating the human sequence of SGCG, under the control of a desmin promoter, at a dose of 2e13 vg/kg. (**b)** Histological characterization of GA muscle. Top panel: representative images of GA cross-sections immunostained with antibody specific for the human SGCG. Bottom panel: representative images of GA cross-sections labeled with HES. (**c)** Western blot analysis of GA muscles for the expression of human SGCG. GAPDH is used as loading control. (**d)** Viral copy number (VCN) quantification in the GA muscle. Data are presented as scatter plot (average ± SD) (n=10, right and left GA muscles are analyzed). (**e)** Relative SGCG mRNA expression in the GA normalized to Rplp0. Scatter plots represent average ± SD. (**f)** Dosage of CK in serum of mice 4-weeks post-injection. Data are presented as scatter plot (average ± SD). (**g)** ELISA quantification of MYOM3 in the serum of the mice 4-weeks post injection. Data are presented scatter plot (average ± SD). (**h)** Global force evaluation by an escape test before sacrifice of mice. Data are presented as scatter plot (average ± SD) of global force normalized to mice body weight. (**i)** Relative Lgals3 mRNA expression in the GA normalized to Rplp0. Scatter plots represent average ± SD. (**j)** Quantification of double positive puncta (LAMP2+LGALS3+) in the myofibers of GA muscles. Data are represented as scatter plots with median (n>338). (**k)** Representative confocal images of transversal sections of GA muscle immunostained for LAMP2 (green) and LGALS3 (red). An ANOVA test with Tukey multiple comparison was used for all statistical comparisons. ANOVA p-value *p<0.05, **p<0.05, ***p<0.001, ****p<0.0001, ns=non-significant.

**Fig. S9.**
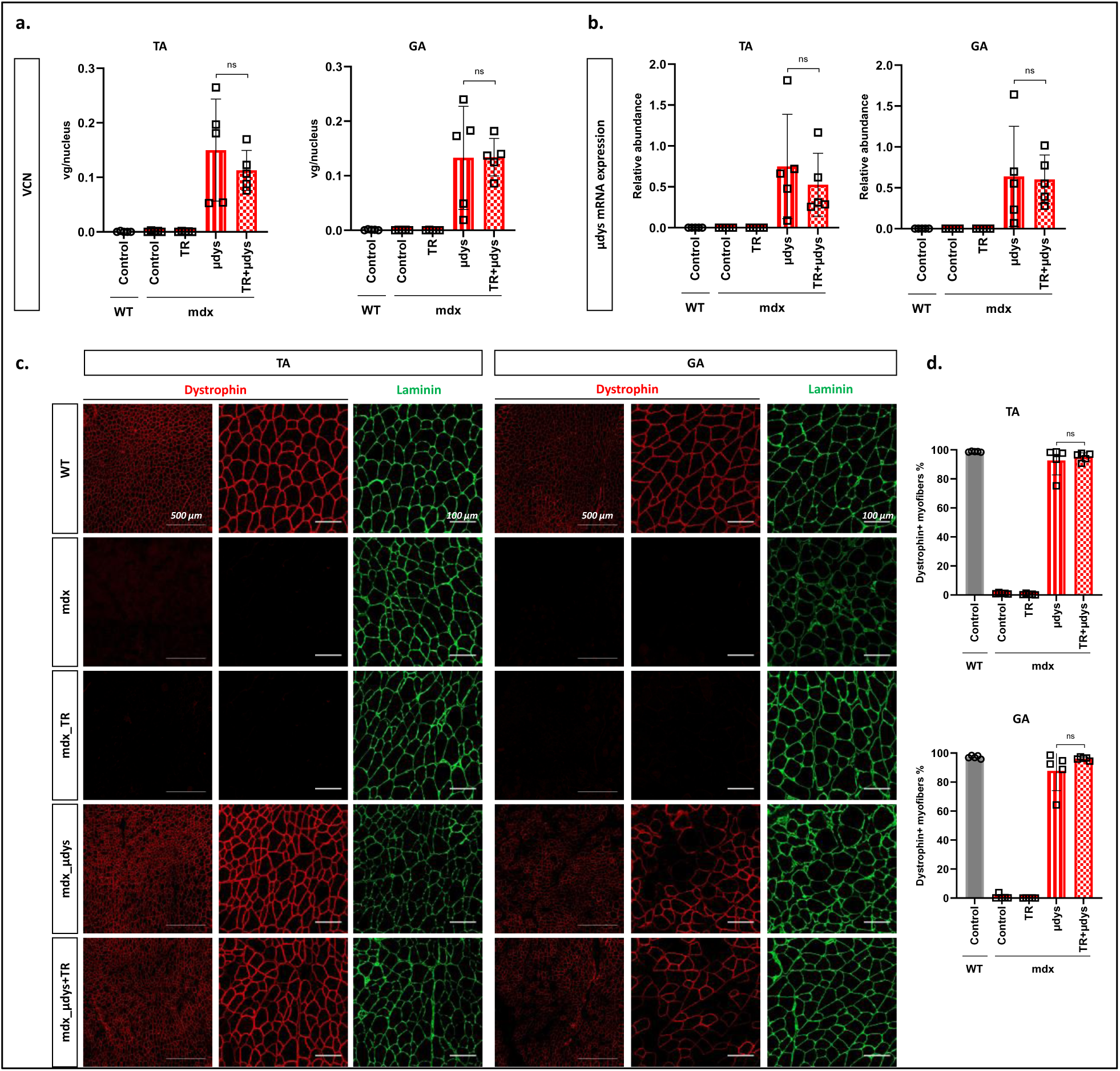
Trehalose does not affect rAAV infectivity or transgene expression. **(a)** Viral copy number (VCN) quantification in the TA and GA muscles. Data are presented as scatter plot (average ± SD). (**b)** Relative µdys mRNA expression in the TA and GA muscles, normalized to Rplp0. Scatter plots represent average ± SD. (**c)** Representative fluorescence images of transversal sections of TA and GA muscles immunostained for Dystrophin (N-terminal) and Laminin. (**d)** Quantification of Dystrophin positive fibers in the TA and GA muscles. Data are presented as scatter plots (average ± SD). An ANOVA test with Tukey multiple comparison was used for all statistical comparisons. ANOVA p-value ns=non-significant.

**Fig. S10.**
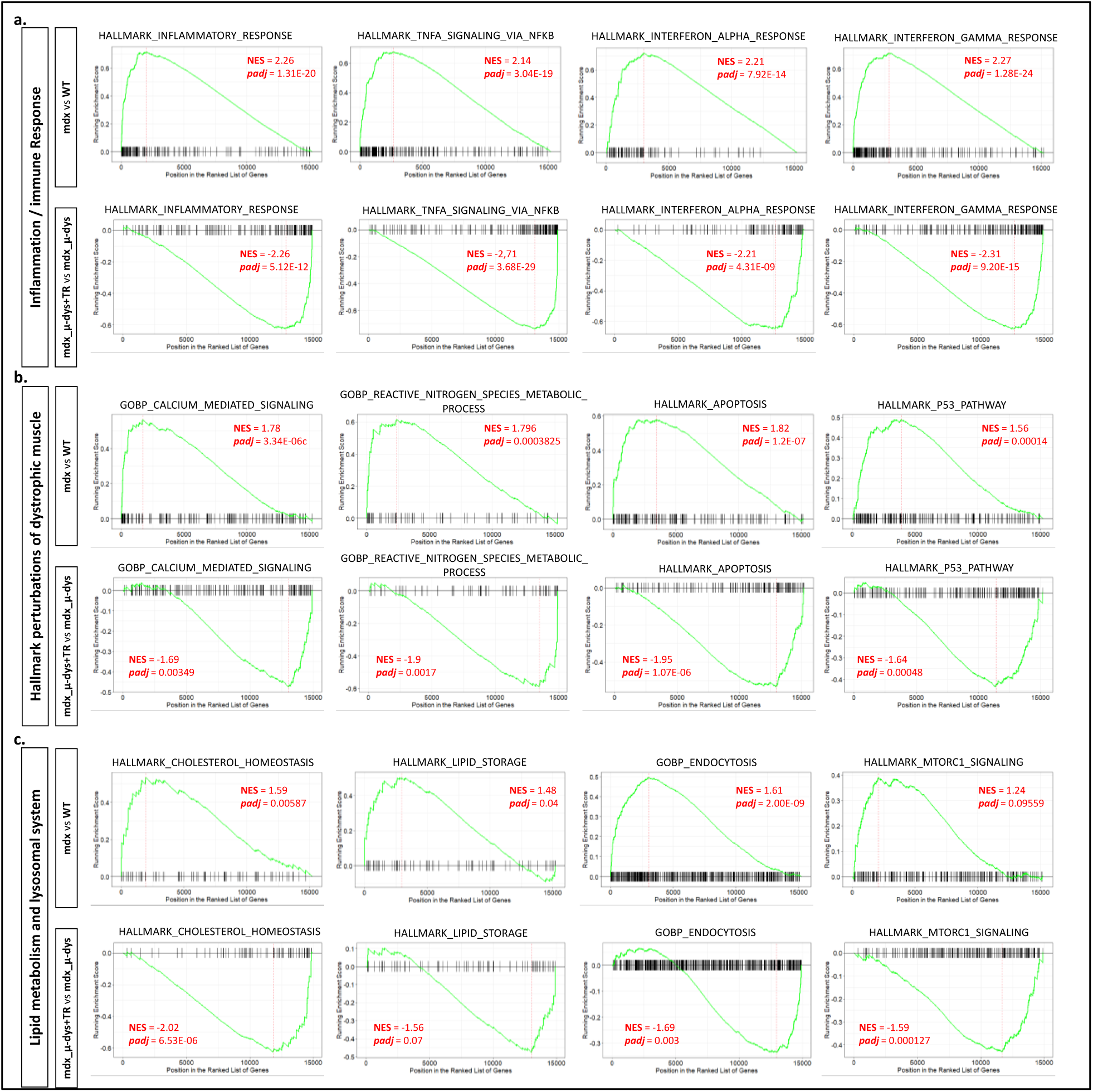
Gene set enrichment analysis (GSEA) of various dysregulated pathways. **(a)** Pathways related to inflammation and immune response. (**b)** Hallmark dysregulated pathways in the dystrophic muscle, including calcium signaling, reactive nitrogen species stress and apoptosis. (**c)** Dysregulated pathways related to lipid metabolism (cholesterol homeostasis, lipid storage) and lysosomal system (endocytosis). MTORC1 signaling is known to act on both lipid metabolism and trafficking, and on the lysosomal system. Upper panels represent a comparison between mdx and WT. Lower panels represent a comparison between µdys only and µdys+trehalose treated groups. NES: Normalized enrichment score, padj: BH-adjusted p-value.

**Table S1.** Patients’ biopsies details.

| **Case number** | **Disease** | **Mutation** | **Age** |
| --- | --- | --- | --- |
| Patient 1 | Dystrophinopathy | DMD del<13-36> | 2 years |
| Patient 2 | Control | - | 2 years |
| Patient 3 | Dystrophinopathy | DMD deletion <53-55> | 3 years 10 months |
| Patient 4 | Control | - | 3.5 years |
| Patient 5 | Sarcoglycanopathy | gSARC mutation homo del525T | 9 years |
| Patient 6 | Control | - | 9 years |

**Table S2.**
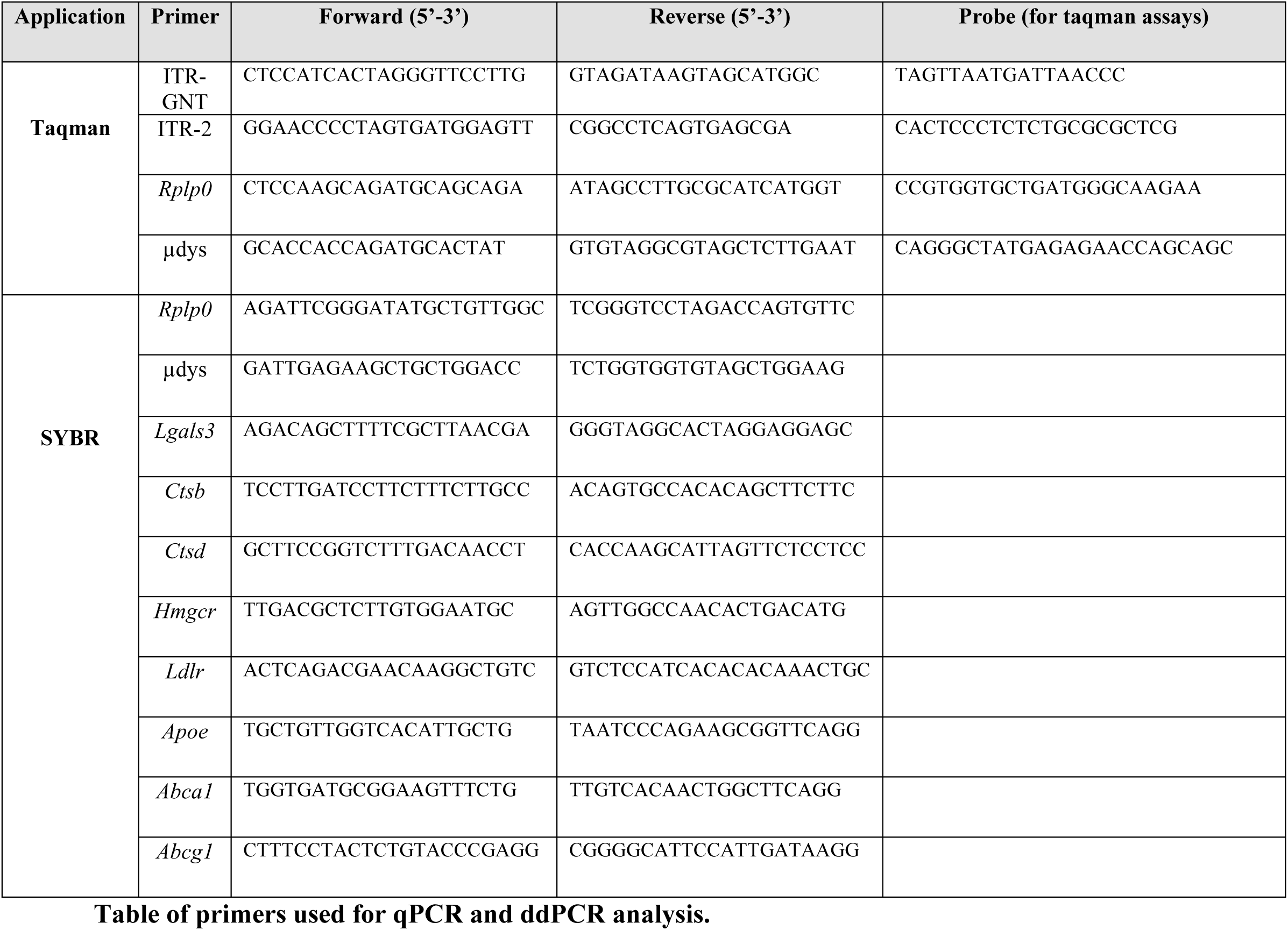
Table of primers used for qPCR and ddPCR analysis.

**Table S3.** Antibodies details for western blotting.

| **Antibody** | **Supplier / Reference** | **Species** | **Dilution** |
| --- | --- | --- | --- |
| Anti-CHMP4B | ProteinTech group, 13683-1-1P | Rabbit Polyclonal | 1:500 |
| Anti-GAPDH | Sigma, G9545 | Rabbit Polyclonal | 1:1000 |
| Anti-GAPDH | Sigma, PLA03023 | Goat Polyclonal | 1:1000 |
| Anti-LAMP2 | Abcam, Ab13524 | Rat monoclonal | 1:1000 |
| Anti-LC3B | Sigma, L7543 | Rabbit Monoclonal | 1:1000 |
| Anti-LGALS3 | R&D, AF1197 | Goat Polyclonal | 1:1000 |
| Anti-SGCG | Abcam, ab203112 | Rabbit Monoclonal | 1:2000 |
| Anti-SQSTM1 | Sigma, P0067 | Rabbit Polyclonal | 1:1000 |
| Anti-ALIX | ProteinTech group, 12422-1-AP | Rabbit Polyclonal | 1:2000 |

**Table S4.**
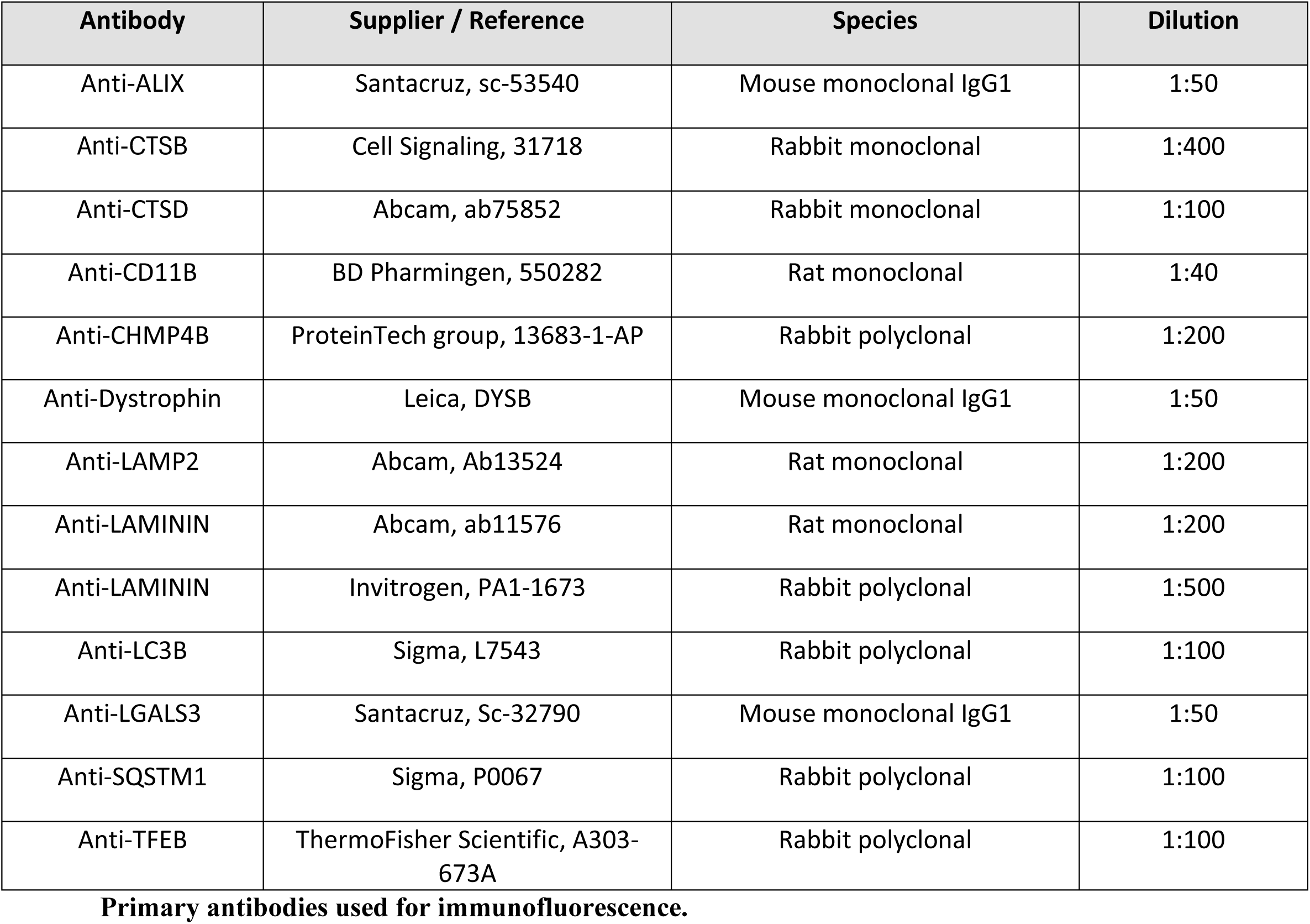
Primary antibodies used for immunofluorescence.

**Table S5.** Secondary Antibodies.

| **Application** | **Antibody** |
| --- | --- |
| **Western Blot** | Licor, 926-68024, Donkey Anti-Goat 680 |
|  | Licor, 926-32214, Donkey Anti-Goat 800 |
|  | Licor, 926-68073, Donkey Anti-Rabbit 680 |
|  | Licor, 926-32213, Donkey Anti-Rabbit 800 |
|  | Licor, 926-32219, Donkey Anti-Rat 800 |
| **Immunofluorescence** | Invitrogen, A11006, Goat-anti Rat 488 |
|  | Invitrogen, A21094, Goat-anti Rat 633 |
|  | Invitrogen, A11005, Goat-anti Mouse A594 |
|  | Invitrogen, A21125, Goat anti-Mouse IgG1 A594 |
|  | Invitrogen, A11008, Goat-anti Rabbit A488 |
|  | Invitrogen, A11037, Goat-anti Rabbit A594 |

## Notes

### Competing Interest Statement

The authors have declared no competing interest.

